# Diffusion and distal linkages govern interchromosomal dynamics during meiotic prophase

**DOI:** 10.1101/2021.04.23.440859

**Authors:** Trent A. C. Newman, Bruno Beltran, James M. McGehee, Daniel Elnatan, Cori K. Cahoon, Michael R. Paddy, Daniel B. Chu, Andrew J. Spakowitz, Sean M. Burgess

## Abstract

The pairing of homologous chromosomes (homologs) in meiosis is essential for distributing the correct numbers of chromosomes into haploid gametes. In budding yeast, pairing depends on the formation of 150-200 Spo11-mediated double-strand breaks (DSBs) that are distributed among 16 homolog pairs, but it is not known if all, or only a subset of these DSBs, contribute to the close juxtaposition of homologs. Having established a system to measure the position of fluorescently tagged chromosomal loci in 3D space over time, we analyzed locus trajectories to determine how frequently, and how long, loci spend colocalized or apart. Continuous imaging revealed highly heterogeneous cell-to-cell behavior of foci, with the majority of cells exhibiting a “mixed” phenotype where foci move into and out of proximity, even at late stages of prophase, suggesting that the axial structures of the synaptonemal complex may be more dynamic than anticipated. The observed plateaus of the mean-squared change in distance (MSCD) between foci informed the development of a biophysical model of two diffusing polymers that captures the loss of centromere linkages as cells enter meiosis, nuclear confinement, and the formation of Spo11-dependent linkages. The predicted number of linkages per chromosome in our theoretical model closely approximates the small number (~2-4) of estimated synapsis-initiation sites, suggesting that excess DSBs have negligible effects on the overall juxtaposition of homologs. These insights into the dynamic interchromosomal behavior displayed during homolog pairing demonstrate the power of combining time-resolved *in vivo* analysis with modeling at the granular level.

**Significance Statement:** Essential for sexual reproduction, meiosis is a specialized cell division required for the production of haploid gametes. Critical to this process is the pairing, recombination, and segregation of homologous chromosomes (homologs). While pairing and recombination are linked, it is not known how many linkages are sufficient to hold homologs in proximity. Here, we reveal that random diffusion and the placement of a small number of linkages are sufficient to establish the apparent “pairing” of homologs. We also show that colocalization between any two loci is more dynamic than anticipated. Our study is the first to provide observations of live interchromosomal dynamics during meiosis and illustrates the power of combining single-cell measurements with theoretical polymer modeling.

---

**D**uring meiosis prophase I, homologous chromosomes undergo pairing, synapsis, and crossing over to ensure their proper segregation at meiosis I. An overarching question is how each chromosome identifies, and pairs with, its homolog partner within, the complex nuclear environment that includes non-homologous chromosomes (1, 2, 3, 4). The general view is that pairing is achieved through homology-based mechanisms that can bring the axes of chromosome pairs into close juxtaposition such that discrete pairing interactions, in conjunction with the establishment of synapsis, are sufficient to align homologs end-to-end (5). While the intermediate steps leading to pairing are not well understood, the process itself is thought to be stochastic with heterogeneity from cell-to-cell.

The budding yeast, *Saccharomyces cerevisiae*, is an important model for the study of homolog pairing as it has been used extensively for characterizing many of the other dynamic events that occur over the course of meiotic prophase I that are now known to be conserved across phyla. These include the transition from the Rabl (centromeres clustered) to bouquet (telomeres clustered) configurations; telomere-led chromosome movement driven by cytoskeletal motor proteins via the LINC complex; the formation and repair of Spo11-induced DNA double-strand breaks (DSBs); and, the assembly and disassembly of the synaptonemal complex (SC): a ribbon-like structure that joins homologs together along their lengths (Fig. 1) (5, 6, 7, 8, 9, 10, 11). Several theoretical models of pairing in yeast have been developed that take into account chromosome size, linkage numbers, and the attachment and motion of telomeres at the nuclear envelope (12, 13, 14, 15, 16), yet no study to date has combined biophysical modeling together with mpirical measurements of meiotic “pairing” dynamics in live cells.

**Fig. 1.**
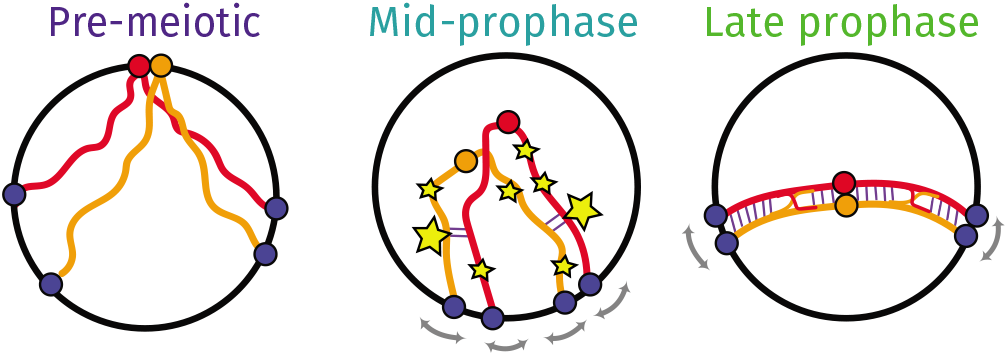
Overview of chromosome conformations in pre-meiotic cells (*T_M_* = *T*_0_) and in meiotic cells in mid-prophase (~ *T*_3_, *T*_4_) and late prophase (~ *T*_5_, *T*_6_). At *T*_0_ cells are in the G0 stage prior to DNA replication and chromosomes are arranged in the Rabl configuration with centromeres clustered at the nuclear periphery (56). Following transfer to sporulation media, the meiotic program begins with cells entering S-phase, over which time the centromeres are dispersed and telomeres start to cluster in the bouquet (56, 57, 58, 59, 60). At early- to mid-prophase, Spo11 initiates the formation of DSBs (61), shown as stars, of which the majority are repaired using the homologous chromosome as a substrate (9) (homologs are red and orange lines; note that each lines in mid- and late-prophase represents the pair of newly replicated sister chromatids). DSBs that go on to form Class I or “interfering” crossovers, shown as the large stars, assemble the synapsis initiation complex (SIC) (33, 41, 42), where the new SC is shown as blue lines. Concomitantly, telomeres are subject to motion driven by cytoskeletal motor proteins shown as gray arrows (7, 62). By late prophase, homologs are synapsed end-to-end and with crossover intermediates maturing into crossover products as shown.

Homolog pairing in yeast has been studied using a number of different assays including fluorescence *in situ* hybridization applied to spread chromosome preparations (17, 18), a “collision” assay based on *Cre*/*loxP* recombination measuring the relative position and accessibility of pairs of homologous loci (19), and a fluorescence reporter operator system (FROS) that enables specific chromosomal loci to be tagged and followed microscopically in live cells. When allelic loci on homologous chromosomes are tagged, this “one-spot, two-spot” assay has been used as a proxy for local homolog juxtaposition (Fig. 2) (2, 4, 20, 21, 22, 23, 24, 25, 26). However, with only a static snapshot, it is not possible to know if colocalization represents a true homolog pairing interaction; that is, if the foci remain colocalized until homologs are segregated at anaphase. While it has been proposed that homologs may undergo many transient interactions that become progressively stabilized throughout prophase (27), this has not yet been investigated.

**Fig. 2.**
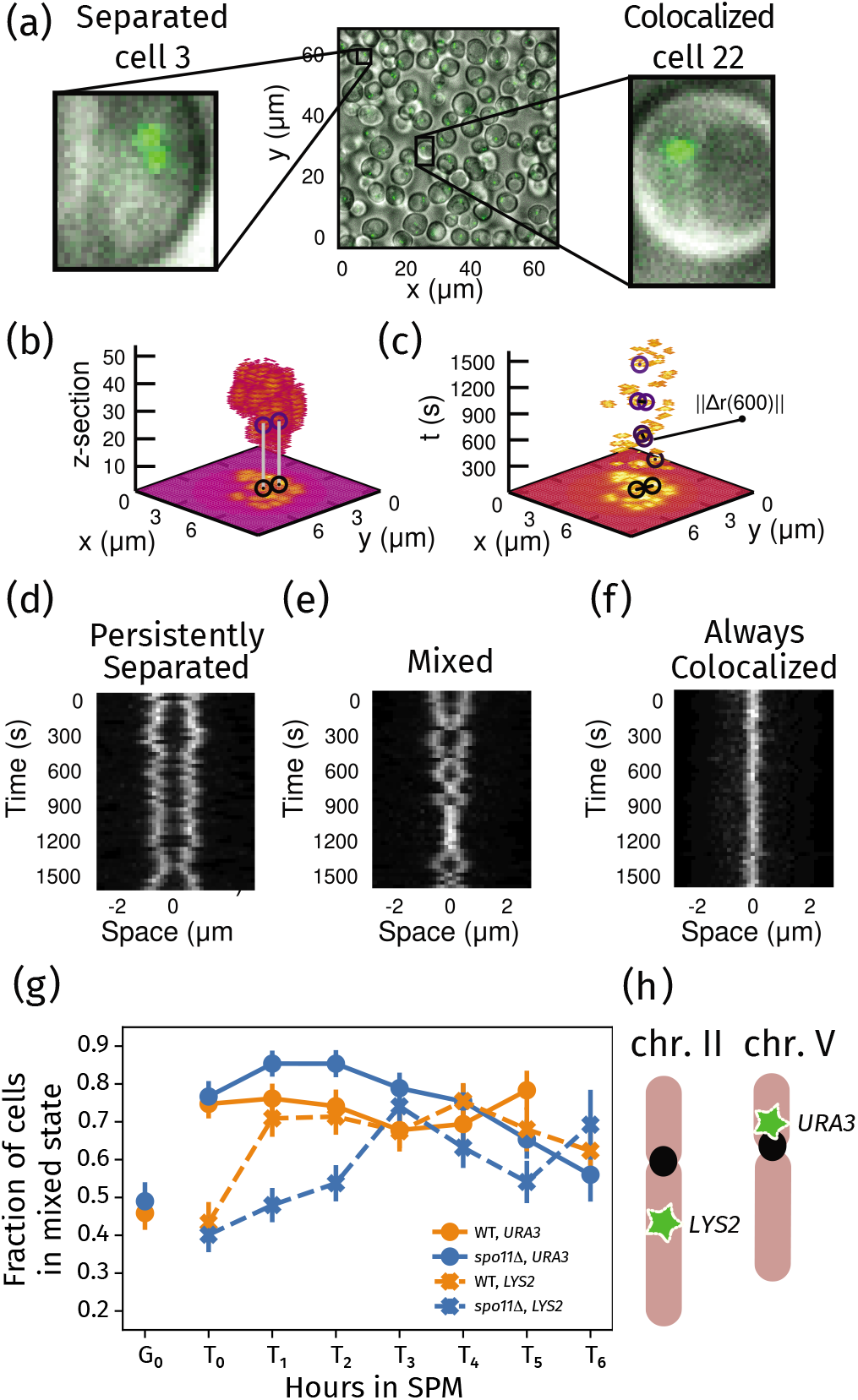
(a) A typical field of cells, highlighting example cells showing either two spots (left) or one spot (right). (b-c) Maximum intensity projections (MIPs) of the relative positions of fluorescent foci at 30 s intervals. In (b), the vertical axis corresponds to a z-stack (with step size 0.25 μm). For each x and y coordinate, the maximum value over all time points for that z-stack is shown. In (c), the vertical axis represents time (*t*, in seconds), and the projection is instead performed over z-stacks. The positions of the loci and the distance between them is highlighted for select time points. (d-f) kymographs showing the distance between the loci in a single cell over the 25 minute imaging period. Each horizontal slice in the kymograph shows the fluorescence intensity along the line joining the centers of the two loci in a single frame. Example of cells where the loci are separated (d), or colocalize (f) for every frame. The “mixed” cell shown in (e) undergoes several transitions between the two states. (g) Fraction of cells in the mixed state versus hours in SPM through meiosis for the *URA3* and *LYS2* loci in wild-type and *spo11*Δ. cells. Plot was made from aggregating all available data for each meiotic stage. The error is the standard error of the mean with the sample count set to the number of trajectories (see Supplemental File 2). The error is the standard error of the mean with the sample count set to the number of trajectories. (h) Schematic representation of the genomic positions of the *URA3* and *LYS2* loci on chromosomes V and II, respectively.

Though the mechanisms promoting homolog colocalization are not well understood, in yeast inter-homolog linkages depend on the formation and repair of DSBs created by Spo11 and its partners during prophase I (9). For any given cell in meiosis, any sequence has the “potential” (albeit not all equally) to experience a DSB. While 150-200 DBSs are formed per cell, only ~90-94 DSBs go on to form crossovers (CO). Another ~66 are repaired using the homologous chromosome but do not lead to CO formation, called non-crossovers (NCOs), and the remaining are repaired with the sister chromatid (28, 29, 30, 31, 32). COs are divided into Class I and Class II. Class I COs account for ~70% of total COs; their position and number are specified in mid prophase by the ZMM proteins that make up the synapsis initiation complex (SIC), which functions to couple homologous recombination with the establishment of the SC (8, 33, 34, 35, 36, 37, 38, 39, 40, 41, 42, 43). Class II crossovers arise from an alternative repair process that does not involve the SIC and are “interference independent” (44, 45, 46). Thus, the question arises; are the excess DSBs necessary to mediate pairing, or is the smaller number that go on to form crossovers (Class I and/or Class II) sufficient?

Rather than the homolog pairing process being independent for each “paired” locus, several models relating meiotic homolog pairing to polymer theory predict that “pairing” at one locus will increase the probability that pairing at an adjacent site will occur (14, 15, 16, 47, 48). That is, a molecular linkage at one site on the chromosome is expected to restrict the diffusive properties of adjacent sites along that chromosome (49, 50, 51). However, this has not been explicitly evaluated experimentally in the case of meiotic homolog pairing. Furthermore, it is not known if the repair of Spo11 DSBs leads to any directed motion that could aid in bringing homolog axes into close juxtaposition, similar to the observed DSB-dependent directed motion that brings telomeres into proximity seen in ALT cells (Alternative Lengthening of Telomeres) (52). For instance, it has been proposed that single stranded DNA filaments, formed by resection of DSBs, might capture a locus of the homologous chromosome and processively “reel” the axis into alignment (53, 54, 55).

To address these gaps in knowledge, we observed the behavior of FROS-tagged loci in 3D space over time, and show, for the first time, the highly dynamic behavior between loci on homologous chromosomes during meiosis prophase I. In contrast to static snapshots, continuous imaging revealed that the majority of cells show a “mixed” phenotype in which foci alternate between colocalized and separated states, indicating that once “paired”, homologous loci need not remain paired until anaphase. We then used our experimental measurements of the dynamic changes in distance between homologous loci to develop a theoretical model of homolog pairing based on polymer diffusion in the viscoelastic medium of the nucleus. This modeling suggests that homolog pairing is a fluctuation-driven process, rather than the result of chromosomes being pushed or pulled together via an active mechanism. Moreover, the addition of a a small number of link-ages (between 2 and 4) per chromosome pair, closely approximating the number of Class I crossovers, accounts for the observed level of confinement, while the position of linkages and other factors account for the heterogeneous cell-to-cell behavior. These insights illustrate the utility of combining live imaging with biophysical modelling for the study of dynamic processes in living cells.

## Results

### Homologous interactions remain transient throughout meiosis

Our study used yeast strains carrying chromosomally-integrated *tet* operator arrays of 112 repeats bound by fluorescent TetR-GFP protein (25). Operators were inserted at either the *URA3* locus—which is on the short arm of chr. V (577 kb) near the centromere, or the *LYS2* locus—which is in the center of the long arm of chr. II (813 kb; see Fig. 2 and Table S1).

Cells were cultured for synchronized progression through meiotic prophase as described previously (63). Briefly, cells were grown in yeast peptone (YP) media containing acetate for arrest in G0. There after, cells were transferred to nutrient-deficient sporulation medium (SPM) where they then undergo S-phase followed by the entry into meiosis prophase I. Aliquots of cells were removed from the culture every hour (*T_M_* = *T*_0_, *T*_1_,…) and imaged over a 25 minute period at 30 second intervals (*t_i_* = 0, 30,…, 1500). Following extensive quality control (see Supplementary Information, Fig S15 and S16), the positions 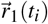 and 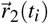 of the two fluorescent foci (or the single focus representing colocalized loci) was determined as seen in Fig. 2.

Our single-cell measurements permitted us to evaluate the kinetics of transient colocalizations between loci based on their XYZ coordinates from over 1.4 million 2D images. In Fig. 3, we report the fraction of time the two loci existed in a colocalized state, defined as foci less than 250nm apart (i.e. their point-spread functions are not distinguishable with a separation of less than 250nm), averaged over all cells imaged across all time courses for each strain and over all frames of each movie. In *spo11*Δ mutants (both for the *URA3* and *LYS2* loci), the fraction of time colocalized continued to decrease over time. However, the wild-type cells exhibited a non-monotonic trend in the fraction of time colocalized (in this case, decreasing then increasing), as the loci spent more time together during the mid- and late-stages of prophase I (times *T*_3_ to *T*_6_). The fraction of time colocalized continued to increase through the late stages (but never reached 100%). Previous studies reporting the fraction of cells with colocalized foci for any given time point, e.g. (25), were unable to distinguish between an increased frequency of transient colocalization on the one hand and the formation of stable “pairing” interactions.

**Fig. 3.**
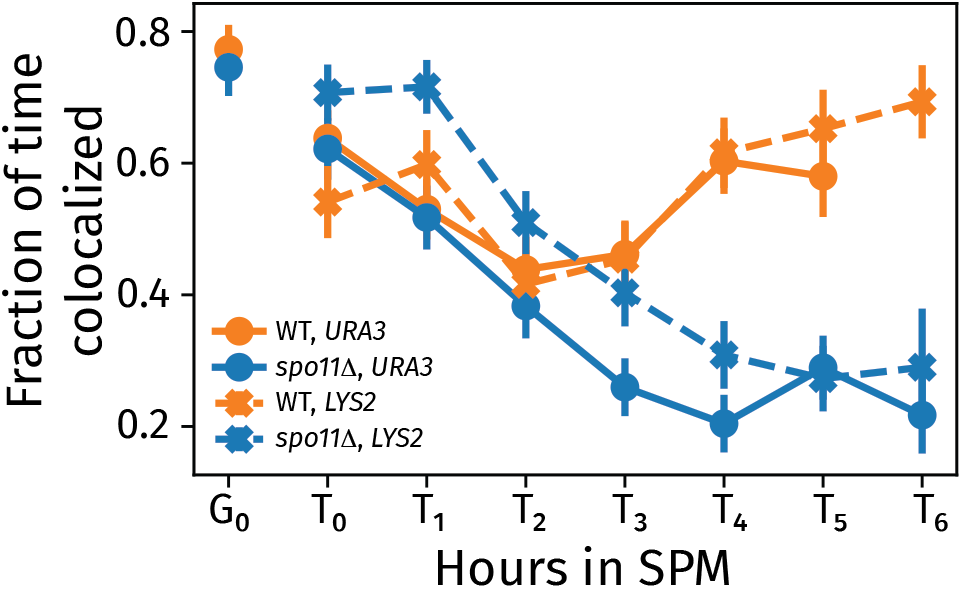
The fraction of time at each stage of meiosis (*T_M_* = *T*_0_,*T*_1_,…) that foci are in a colocalized state for each of the two loci and strains examined. Plot was made from aggregating all available data for each meiotic stage. The error is the standard error of the mean with the sample count set to the number of trajectories (see Supplemental File 2).

Using the dynamic information in our measurements, we classified entire trajectories for individual cells as being persistently colocalized and persistently separated, where the observed state (i.e. colocalized or separated) did not change for the duration of the movie. We also identified a third category of cells with “mixed” trajectories, where the cell was observed to transition into or out of a colocalized state during the 25 minute period. These three states could be distinguished in locus-separation kymographs (see Fig. 2, and Supplementary Information Fig. S2 and Fig. S18). We were surprised to find that the majority of cells were classified as “mixed” suggesting that in the many instances where foci were colocalized, the loci themselves were not necessarily paired.

From cells with “mixed” trajectories, we determined the distribution of dwell times in the colocalized and separated states. Figure 4 shows the probability density function of dwell times for loci in the colocalized state for the *URA3* locus (see Supplementary Information for Fig. S3 for colocalized and separated states and Fig. S4 for corresponding plots for *LYS2*). This data demonstrates the transient nature of the colocalization of the loci throughout the observation period for both the wild-type and *spo11*Δ strains. These plots, shown on a log-log scale, demonstrate the power-law nature of the dwell time distributions.

**Fig. 4.**
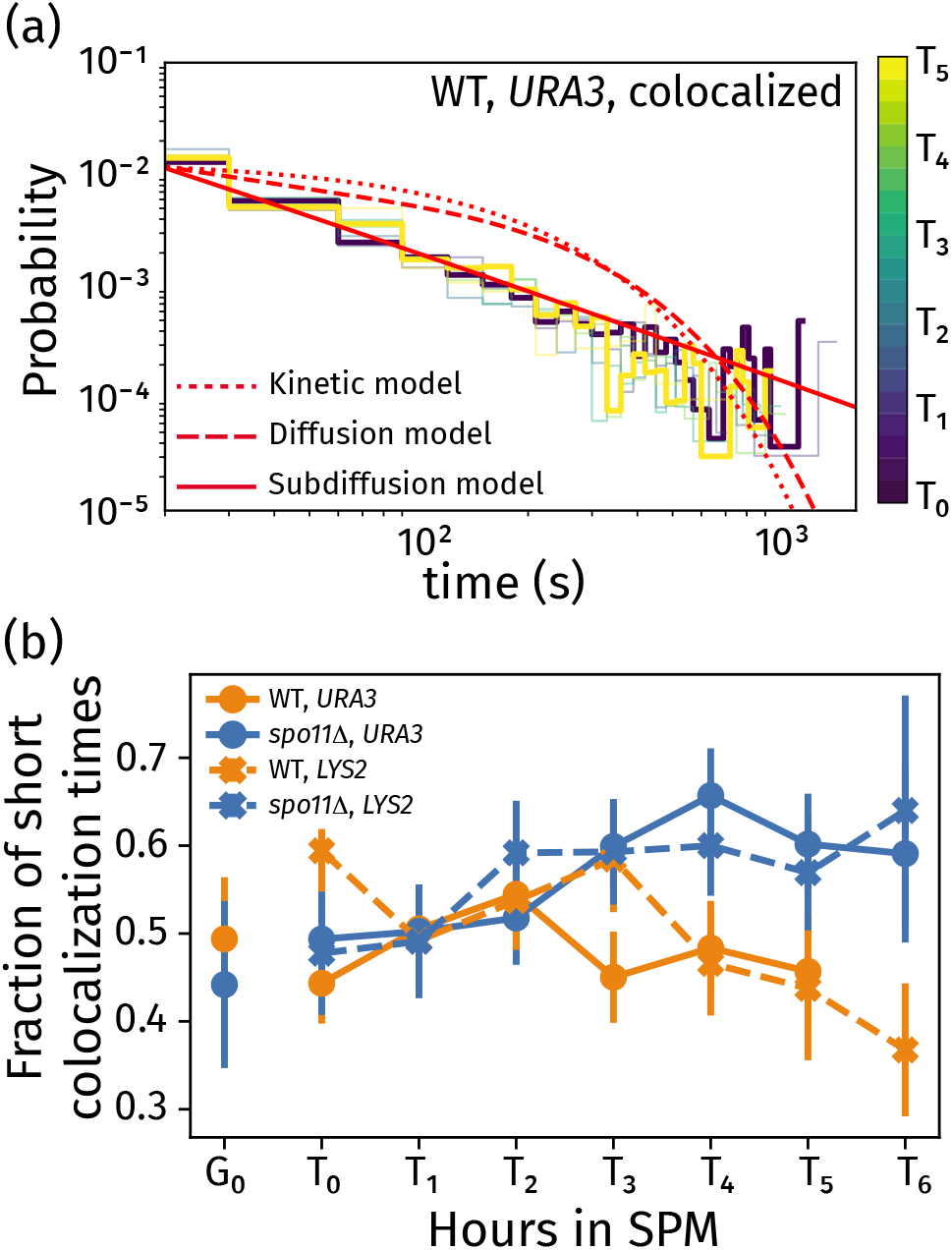
Histograms of dwell times in the colocalized states for the *URA3* locus (a), colored by the time since transfer to sporulation media. Along with the experimental data, we show theoretical fits for kinetic (dotted), diffusion (dashed), and subdiffusion (solid) models. The fraction of short colocalization times (b) gives the probability of colocalization time being less than 30 seconds versus time in sporulation media, including data for wild-type and *spo11*Δ strains for the *URA3* and *LYS2* loci.

We include theoretical curves for three candidate models whose dwell-time distributions are limited by kinetics (dotted curve), diffusion (dashed curve), and subdiffusion (solid curve). A detailed derivation of these three models is provided in Supplementary Information. The kinetic model is governed by an exponential distribution for Poisson-distributed times for transitioning between colocalized and separated states. The diffusion and subdiffusion models are derived from a generalized diffusion process in 1 dimension, representing a prediction for the dwell-time distribution arising from stochastic trajectories. The subdiffusion model coincides with stochastic motion of loci with a mean-square displacement that scales as ~ *t^B^*, where *B* = 0.24. We demonstrate in our subsequent analyses that this specific choice of power-law scaling is consistent with single-locus trajectories within our data.

The experimental measurements exhibited power-law dwell-time distributions, which are more consistent with the subdiffusion model than either the kinetic model or the diffusion model. This indicates that the dwell-time distributions are limited by individual trajectories exhibiting subdiffusive scaling. Such power-law distributions arise in subdiffusion-limited intra-chain processes between polymers due to the multi-scale relaxation of elastic stresses that cause the polymer segments to move subdiffusively (51, 64).

While the general trends in the dwell-time distributions are similar for wild-type and *spo11*Δ strains (see Supplementary Information, Fig. S3 for *URA3* and Fig. S4 for *LYS2*), we note several important distinctions. The colocalization dwell-time distribution for wild-type cells (Fig. 4a) exhibits a marked progression through meiosis (from *T*_0_ in purple to *T*_5_ in yellow) towards favoring longer dwell times in the colocalized state, marked by a long-time tail in the distribution for *T*_5_. This trend is apparent as a reduced fraction of short colocalization times (i.e. the probability for times less than 30 seconds) over the course of meiosis at the *LYS2* and *URA3* loci in wild-type cells (Fig. 4b). In contrast, *spo11*Δ cells (data shown in Fig. S3 and S4) and cells with tags on heterologous chromosomes (labeled “Het”, see Supplementary Information, Fig. S5) showed a higher fraction of short dwell times later in meiosis (Fig. 4b).

### Live imaging reveals physical tethering between homologous loci

Since Spo11-dependent colocalization of homologous loci is evident after 3 hours post transfer to sporulation media (Fig. 4), we expected that trajectories measured in cells after time point *T*_3_ would be influenced by tethering mediated by homologous recombination. This was tested by comparing the maximum value of the mean-square displacement (MSD) curves of individual loci to the mean-square change in distance (MSCD) curves of those same loci. Following (65), we define the MSCD to be the mean-squared change of the vector connecting the two loci, 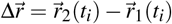. For un-linked loci, we would expect the MSD and MSCD curves to plateau to a comparable value (approximately the square of the confinement radius), since the only source of confinement is the nuclear radius if the chromosomes are not linked. Therefore, a MSCD curve which plateaus to a lower level than the MSD curve is indicative of some level of linkage between the two loci. Supplementary Information (Fig. S7) provides the comparison between MSD and MSCD for the *URA3* and *LYS2* loci, confirming the MSCD curves are substantially smaller than the MSD values. We note that MSD is more susceptible to artifacts during the image collections (e.g. stage drift) or cellular motion (e.g. nuclear rotation) than MSCD. Our analyses limit the impact of drift by excluding sections of movies where excessive drift occurred.

Figure 5 shows time-averaged, single-cell MSCDs for a random subsample of cells from a single movie of *URA3* at *T*_5_. We computed the time average for a single trajectory as

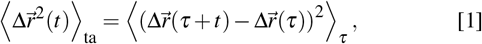

where 〈·〉_*τ*_ indicates the averaging is performed over all possible values of *τ*. Time stamps where two foci could not be resolved were omitted from all MSCD calculations, meaning that we are explicitly computing the dynamics from movie frames where the loci are non-overlapping (we test for any bias resulting from this element of the experimentation below). The Supplementary Information (Fig. S8 for *URA3* and Fig. S9 for *LYS2*) provides plots of the single-cell MSCDs for times *T*_0_ to *T*_5_ for wild-type and *spo11*Δ strains.

**Fig. 5.**
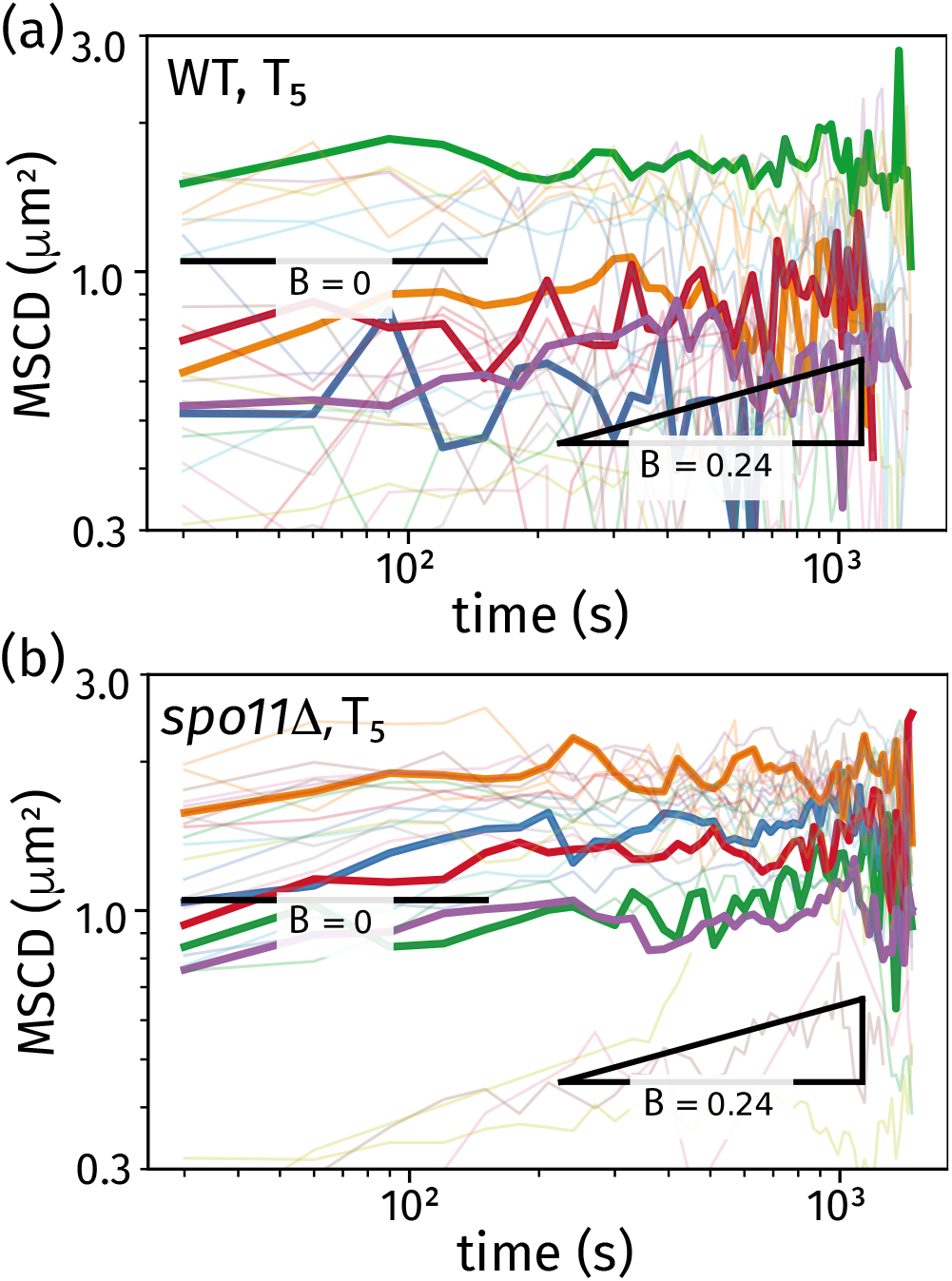
Single-cell MSCDs for *URA3* trajectories at *T*_5_. These plots show results from 25 randomly selected cells (light) along with 5 randomly selected cells (bold) for wild-type cells (a) and *spo11*Δ cells (b). Each plot includes two power-law scaling behaviors associated with confined motion (slope *B* = 0) and unconfined subdiffusive polymer motion (slope *B* = 0.24).

Figure 5 shows results of the behavior of 25 randomly selected cells (light color) from the wild-type population (top), and the *spo11*Δ population (bottom) from which we highlight data from 5 cells to demonstrate the cell-to-cell heterogeneity and individual-cell behaviors. The trajectories exhibit a combination of power-law transport (MSCD ~ *t^B^*) and confined motion (constant MSCD), indicative of initial subdiffusive transport followed by spatial confinement at a plateau value. This behavior is also seen at *T*_0_ for both *URA3* and *LYS2* loci in the wild-type strain and is reported in the Supplementary Information (Figs. S10 and S11). This analysis includes a fit of each single-cell MSCD to a function *MSCD* = min(*At^B^*,*C*), which exhibits an initial power-law behavior followed by a plateau. This analysis identifies the scaling coefficient *B* for individual trajectories, and we constructed a histogram of *B* values for those trajectories with at least 10 data points in the power-law regime. From this analysis, the distribution of values of the power-law slope *B* ranged from about zero to 0.5, with an average value of *B* = 0.24. Figure 5 shows the power-law scaling behaviors associated with confined motion (zero slope) and the experimentally determined power-law scaling (slope *B* = 0.24) as guides.

The MSCD behaviors of wild-type (a) and *spo11*Δ (b) at *T*_5_ showed distinct differences that reflect their underlying biological states. At this late stage of prophase I, we anticipate that most cells are no longer in the Rabl configuration and homologous chromosomes have paired via Spo11-dependent recombination interactions (56, 57, 59). The *spo11*Δ cells show a clustering of the MSCD plateau between 1 *μ*m^2^ and 2 *μ*m^2^, which we associate with confined motion within the nuclear environment. Notably, several individual cells in Fig. 5b exhibited a significantly lower MSCD plateau, which is likely due to the less-frequent cases of cells remaining in the Rabl configuration at *T*_5_ or cells where centromeres are attached to spindle fibers and about to go through anaphase (56, 66). The wild-type cells in Fig. 5a showed a larger degree of heterogeneity in MSCD behavior.

In summary, the observed heterogeneity in the MSCD plateaus over long time scales indicates three contributions to confinement. (1) the physical boundaries of the nucleus, (2) centromere linkage for cells in the Rabl configuration, and (3) linkages between the homologous chromosomes as prophase I progresses.

### Tethering of homologous loci through random linkages can recreate the range of confinement observed experimentally

To predict the impact of these three sources of confinement on chromosome motion during prophase I, we developed a theoretical model that describes the experimentally observed behaviors. Chromosomal behaviors in living cells, including bacteria (67, 68, 69), mammalian cells (70), and yeast nuclei (16, 69, 71, 72), can be captured by polymer-physics models. These works are generally based on the Rouse model (51) in which the polymer chain is represented as a linear chain of beads connected by springs and motion is driven by random Brownian forces. Several treatments of *in vivo* chromosomal dynamics extend the Rouse model to include the influence of viscoelasticity, which we refer to as the viscoelastic Rouse model (67, 68, 69, 70). Adding viscoelastic stresses to the polymer dynamics leads to a significant reduction in the power-law scaling of various metrics (e.g. MSD, MSCD, and the velocity autocorrelation function) (49, 68).

The original Rouse model exhibits a monomer MSD with power-law scaling of *t*^1/2^. On the other hand, the viscoelastic Rouse model, for a fluid with scaling exponent *α* (i.e. particle motion exhibits MSD ~ *t^α^*), predicts a monomer MSD with scaling MSCD ~ *t*^*α*/2^. Given the average power-law scaling for our experimental MSCDs having a *B* = 0.24, our results are consistent with a viscoelastic Rouse model with *α* = 2*B* = 0.48. Previous measurements of chromosomal motion in living cells result in *α* values ranging from *α* = 0.78 in bacteria (67, 68, 69) to *α* = 1 in yeast (70, 73) and mammalian cells (16, 69, 71, 72). The lower *α* value derived from our analysis may indicate elastic properties within the viscoelastic environment of the nucleus that were not previously observed (note, *α* = 1 is purely viscous, and *α* = 0 is purely elastic).

We developed a polymer-physics model of homologous chromosomes that extends the viscoelastic Rouse polymer by adding several key physical contributions. First, we confine two Rouse polymers within a sphere of radius *a*, representing the observed nuclear confinement. Second, we link the centromere position of the two polymers (chosen approprtately for the specific chromosome being modeled) to the nuclear envelope, when the cell is in the Rabl configuration (i.e. in G0). Third, we model the effects of homologous recombination by adding linkages between the two polymers with an increasing average number as cells transit through prophase. Our model therefore has the following physical parameters: the Kuhn length *b* of the polymer chains, the spherical radius *a*, the rate constant for transitioning from the Rabl configuration *k*_rabl_, the average number of linkages *μ* (which varies with hours in SPM), and the subdiffusion constant *D*_0_ for polymer segmental motion. The polymer lengths and segmental positions of the tracked loci and centromeres are determined from the genomic properties (Table S1).

Our coarse grained representation enabled us to predict behavior at experimentally relevant time scales (i.e. ranging from seconds to minutes). More detailed molecular models provide additional information regarding local structure dynamics and have addressed chromosomal features including loop extrusion and heterogeneous properties (74), centromere tethering (75), and chromosomal crosslinking (76). Our approach complements these detailed models by predicting large-scale chromosomal dynamics and including influences from some of these detailed effects.

Experimental behavior under various conditions permits us to isolate and determine individual physical parameters in our model Figure 1. First, we assume that the behavior of the MSCD at *T*_0_ (just after induction of sporulation) is dominated by the centromere attachment to the nuclear envelope for the *URA3* locus on chromosome V due to its close proximity to the centromere. We determined the MSCD plateau at time *T*_0_ (MSCD_∞_(*T*_0_)) based on the asymptotic approach to a stable maximum MSCD value. In our model, we assume a linear compaction of the meiotic chromosome to be 31.6 bp/nm; a geometric compaction for a chain with 15 bp linker length (contributing a length of 0.34 nm/bp) and 146 bp compacted into a nucleosome (contributing a length of 0 nm/bp). We then applied our model to chromosome V and calculated the plateau in the MSCD versus Kuhn length (see Supplementary Information, Fig S12) in order to determine the Kuhn length to be *b* = 250nm. We note that previous models of budding yeast chromosomes during interphase predict a linear compaction between 53 and 65 nm/bp and a Kuhn length between 104nm and 170nm (77). If we apply this range of linear compaction to our analysis, our model with equivalent behavior would have a Kuhn length between 121nm and 149nm, which is in the range of reported values for interphase yeast chromosomes (77).

As the cells progress through prophase I, we assume the change in the MSCD of the *spo11*Δ strain arises from progressive transition out of the Rabl configuration. We evaluate the MSCD plateau at each time from *T*_0_ to *T*_6_. We then fitted this data to a function of the form MSCD_∞_ = MSCD_∞_(*T*_0_) exp (–*k*_rabl_*t*) + MSCD_∞_(*T*_∞_) [1 – exp (–*k*_r3bl_*t*)], where *k*_r3bl_ is the rate constant for transition from the Rabl configuration and MSCD_∞_(*T*_∞_) is the MSCD plateau value when the majority of cells have transitioned out of the Rabl configuration. Note, MSCD_∞_ (*T*_0_) is uniquely determined from the *T*_0_ MSCD plateau. From this analysis, we determined *k*_rabl_ = 0.605h^−1^, resulting in an average time for centromere detachment of 1.65 h (between *T*_1_ and *T*_2_). Previous studies have provided data for the fraction of centromere attachment throughout prophase I (56, 59). Our reported *k*_rabl_ value of 0.605h^−1^ is between the values of 0.176h^−1^ (59) and 0.835h^−1^ (56), suggesting that our fitted value is reasonable within the context of previous experiments.

From this fit to the *spo11*Δ data, we extract the theoretical plateau value for the MSCD predicted once all cells have undergone centromere detachment, *MSCD*_∞_(*T*_∞_) = 1.74μm^2^. We then model the MSCD plateau using our theoretical model of two flexible polymers confined within a sphere of radius *a* with their ends attached to the sphere surface (see Supplementary Information for details). Using this model, we determined the best fit sphere radius to be *a* = 1.59 μm. We then used the MSCD plateau values from the wild-type strain for *URA3* to determine the mean number of linkages throughout prophase I to be *μ* = 0.08 at *T*_3_, *μ* = 1.27 at *T*_4_, and *μ* = 3.36 at *T*_5_. We assume the number of linkages between *T*_0_ and *T*_3_ to be negligible, and that the behavior is dominated by centromere attachment during this early stage of prophase I. Similar analyses for the *LYS2* locus yields the mean number of linkages at *T*_5_ to be *μ* = 1.27, and *μ* = 2.06 at *T*_6_ (with *μ* = 0 at earlier times).

We used our theory to test the effect of the imaging resolution of ≈ 250nm on our results. Our experimental analyses of the MSCD neglected values that are less than 0.0625 *μ*m^2^, and we also removed these values from our theory for consistency. Figure S14 provides a comparison between the theoretical curves with and without these excluded values for the *URA3* locus. The quantitative difference between these predictions was negligible for *T*_0_ to *T*_3_ and contributed only a subtle decrease in the predicted MSCD values for *T*_4_ and *T*_5_. This unbiased theory predicts a number of linkages *μ* = 0.08 at *T*_3_, *μ* = 1.20 at *T*_4_, and *μ* = 3.16 at *T*_5_, which has a maximum deviation of 6% from our reported values from the biased theory. Thus, the imaging resolution did not significantly affect the conclusions drawn regarding the phenomena observed in this work.

Figure 6 shows theoretical curves for the MSCD for simulated “cells” (Fig. 6a) that are generated by adding a Poisson-distributed number of “linkage sites” located at random positions along the homologous chromosomes. Figure 6a shows 5 linkage diagrams for simulated “cells”, where the blue lines identify randomly selected linkages. These five “cells” coincide with the five bold MSCD curves in Fig. 6b. In addition, Fig. 6b shows curves for 25 simulated “cells” as light curves (same number of trajectories as presented in Fig. 5), providing a picture of both the individual “cell” behavior and the distribution within the ensemble. Each MSCD curve generated by our theory shows the behavior for a time average over random trajectories for the fixed linkages of each “cell”.

**Fig. 6.**
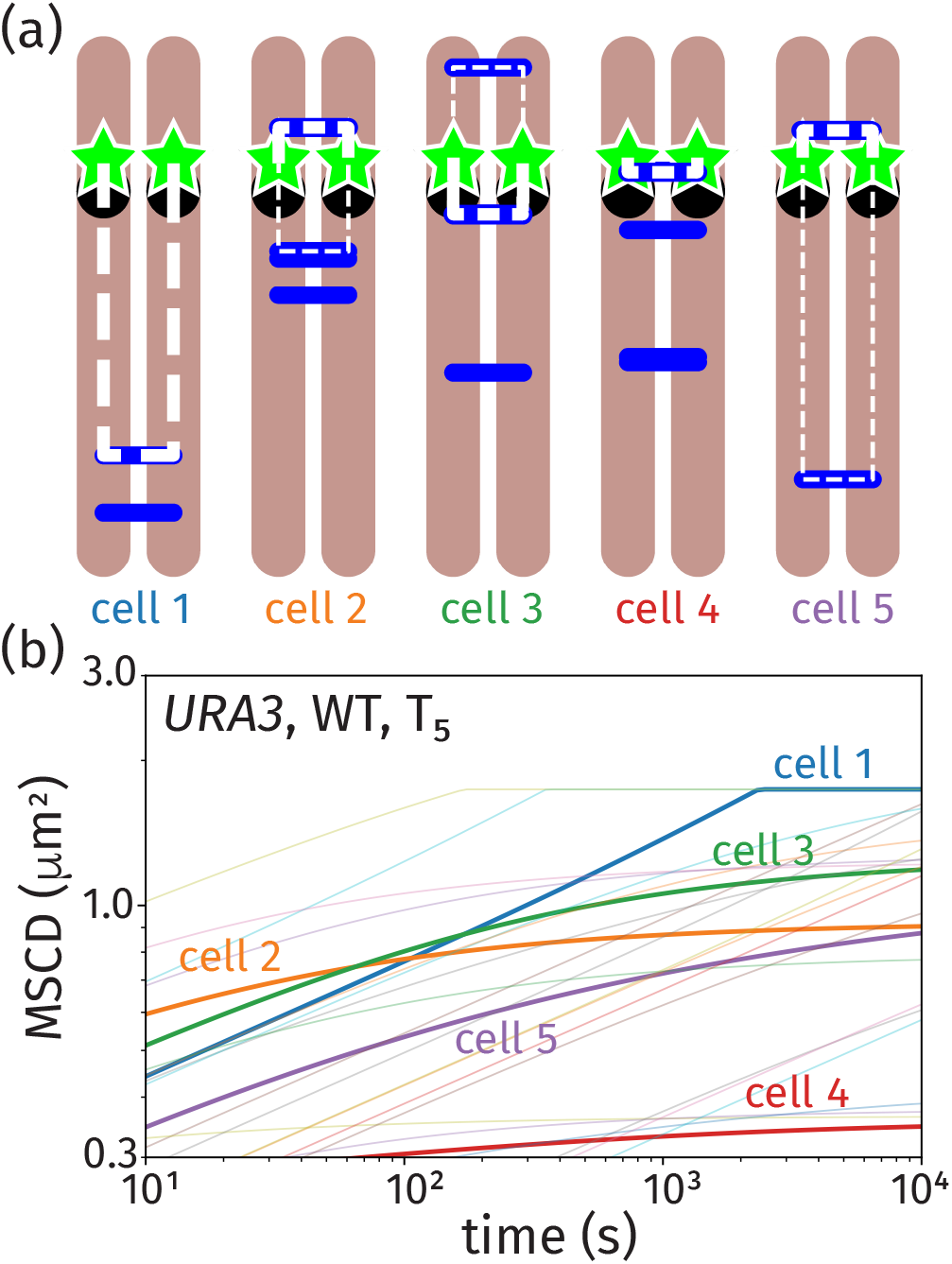
Theoretical curves for the MSCD based on our random-link model for homolog pairing coincident with *URA3* trajectories at *T*_5_. Five individual cell linkage diagrams (a), where the blue sticks identify random linkages along the homologous chromosomes, result in the five bold MSCD curves the plot (b). The MSCD plot shows 25 additional realizations (light) to demonstrate the heterogeneity in the MSCD behavior.

The two copies of our tagged loci behaved as though they were connected by an effective tether whose length is dictated by the distance to the nearest linkage sites, which we highlight in Fig. 6a using bold white for the nearest linkage and thin white for the next-nearest linkage (if applicable). If the tagged locus has a linkage on only one side (e.g. cells 1 and 4 in Fig. 6a), the tagged loci are tethered together by a linear chain. If there are linkage sites on both sides of the tagged locus (e.g. cells 2, 3, and 5 in Fig. 6a), the tagged loci are isolated within an effective “ring” polymer; Figure S1 provides a schematic explanation of the cases where the polymer behaves as a linear versus a ring chain. Assuming these topologies are fixed, we analytically computed the MSCD of the tagged loci by treating them as beads connected by Rouse polymers of appropriate lengths and topology (see Supplementary Information for details on our analytical theory for the MSCD of linear and ring polymers).

The effective tethering radii (MSCD plateau heights) for the randomly linked chromosomes span a similar range as the wild-type data in Fig. 5a. This heterogeneity in behavior arose from variability in the location of the nearest linkage. Instances where a randomly positioned linkage is in close genomic proximity to the tagged locus (e.g. cell 4) resulted in low values of the MSCD plateau. Variability in the distance to the nearest linkage causes the MSCD curves to vary in their magnitude, and there are instances where the nearest linkage is sufficiently far from the tracked locus that it is instead the nuclear confinement that is predicted to dictate the MSCD plateau, as in cell 1 in Fig. 6. Prior to the plateau, each MSCD curve in Fig. 6 exhibits a transient power-law scaling of *t*^0.24^, as dictated by the viscoelastic Rouse model.

### Progression of behavior through prophase I dictated by centromere release and linkage formation

The individual-cell MSCDs at *T*_5_ analyzed above, in Figs. 5 and 6, were useful in determining the tethering effect of interchromosomal linkages at late-stages of meiosis, after a predicted transition from the Rabl configuration. We analyzed the ensemble-averaged MSCD, which pools the behavior of the population of cells, at each meiotic stage (*T_M_*) in order to demonstrate how the biophysical contributions to the dynamics evolve over the course of meiosis. The progression through prophase in our model is marked by two critical events: release of the centromere and formation of Spo11-dependent linkages. The dual time-and-ensemble average MSCD that we used was computed as

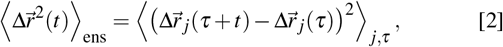

where 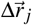 refers to the distance between the two loci in the *j*th cell, and the average is taken over all of the cells in the populations imaged at each *T_M_* (across multiple biological replicates).

In Fig. 7a and Fig. 7b, we show the ensemble-averaged MSCD curves for wild-type and *spo11*Δ strains, respectively, for the *URA3* loci. Fig. 7c and Fig. 7d are the corresponding plots for the *LYS2* loci. From this experimental data, we fitted the subdiffusion coefficients *D*_0_(*T_M_*) at each time using results from our theoretical model, which include Rabl transition and progressive linkage formation (based on analyses from the previous section). The values of the fitted subdiffusion coefficient are provided in the Supplementary Information in Fig. S13. We find that the early-stage data is better fit by a lower diffusivity, and this diffusivity becomes progressively larger as the cells progress through prophase I. Figure 7 shows results of our theoretical model at each time as the solid curves based on 100,000 realizations of our theoretical “cells” whose individual contributions are demonstrated in Fig. 6. The random Brownian motion from each trajectory and cell-to-cell heterogeneity from linkage positioning is smoothed out from the combination of ensemble- and time-averaging within the theory. In our determination of the theoretical average, we excluded MSCD values that are below the detection threshold of 0.0625 μm^2^ to aid comparison with our experimental results that also have this positive bias. The wild-type results in Fig. 7a and c exhibit a non-monotonic behavior (in this case, increasing then decreasing). The *spo11*Δ strain labeled at the *LYS2* locus exhibits monotonic (increasing) behavior in the MSCD (Fig. 7d). The MSCD behavior of the *spo11*Δ strain labeled at the *URA3* (Fig. 7b) exhibits a depressed plateau value at *T*_5_ before increasing again at *T*_6_.

**Fig. 7.**
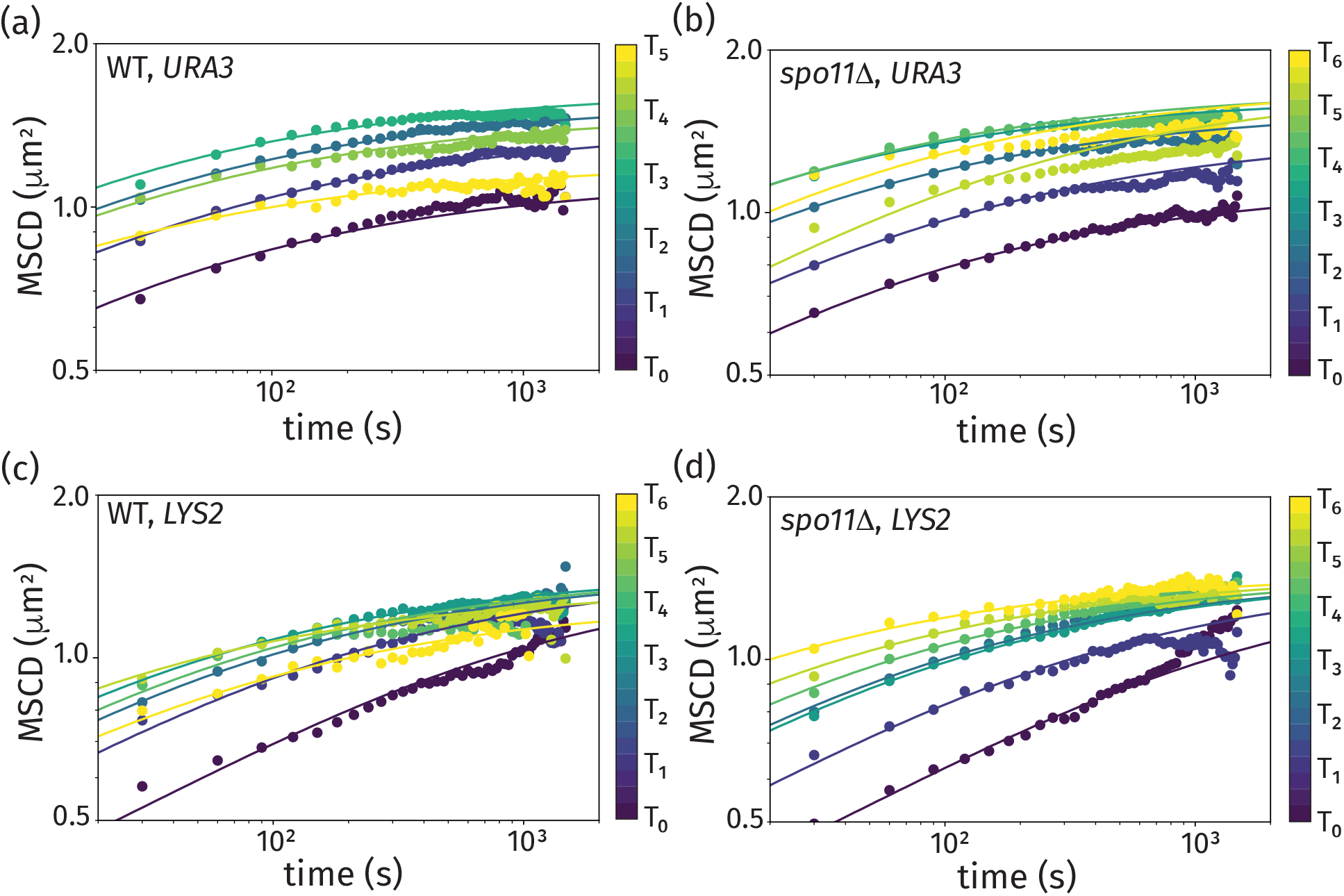
Time-and-ensemble averaged MSCDs at different times after induction of sporulation, for wild-type strain tagged at the *URA3* locus (a), *spo11*Δ strain tagged at the *URA3* locus (b), wild-type strain tagged at the *LYS2* locus (c), and *spo11*Δ strain tagged at the *LYS2* locus (d). Theoretical curves from our model are included for the fitted subdiffusion coefficients.

At early times, the MSCD was substantially reduced due to the effects of two predicted contributors: the large fraction of cells in the Rabl configuration and the reduced subdiffusion coefficient at this early stage. The MSCD increased through this early stage as more cells were no longer linked at the centromere and the subdiffusion coefficient progressively increased. This gradual increase in the subdiffusion coefficient is consistent with previous work (58) that reports significant heterogeneity in the time between induction of sporutation and entry into meiosis, despite the use of synchronized cell cultures. Dissociation of the centromeres from the nuclear envelope is expected to result in an increased plateau of MSCD levels, which was observed. Furthermore, the increases in the subdiffusion coefficient at mid prophase (e.g. at *T*_3_) are consistent with the onset of telomere-led chromosome movements at equivalent time points (78, 79, 80). Notably, the increase in the subdiffusion coefficient is more dramatic for the *URA3* locus than the *LYS2* locus (see Supplementary Information, Fig. S13), which may be due to the closer proximity of the *URA3* locus to a telomere on chr. V (116 kb; 14.8 Kuhn lengths) than the corresponding distance to a telomere for *LYS2* on chr. II (343 kb; 43.3 Kuhn lengths). We hypothesize that genomic proximity to the telomere leads to more efficient physical stress communication; this would result in the rapid prophase movement contributing more significantly to the *URA3* locus than the *LYS2* locus. Further analysis of the impact of rapid prophase movement on locus motion will be explored in future work.

At *T*_3_, when we expect DSBs have begun to form (see Fig.1) (43), the average confinement radius for the *URA3* locus begins to decrease again (see Fig. 7a). Similar behavior was seen for the *LYS2* locus in Fig. 7c, but the inversion was first quantifiable at *T*_4_. In both cases, the decreased MSCD plateaus are consistent with the formation of more linkages between the homologous chromosomes at this time. This reduction in the MSCD plateaus is only expected in wild-type cells, as the *spoii*Δ mutants do not form linkages arising from Spo11-induced double-strand breaks; this was generally true in our experimental data in Figs. 7b and d. However, at time *T*_5_ the *spo11*Δ mutant exhibited a reduced MSCD for the *URA3* locus before returning to the terminal MSCD plateau at time *T*_6_ (see Fig. 7b).

To verify that the observed behaviors in Fig. 7 were specific to homologous chromosomes and not simply due to *spoii*Δ-dependent nuclear compaction, we repeated our analyses in a strain where the FROS tag was integrated in only one homolog of chromosomes V and II at the *URA3* and *LYS2* loci (defined as “Het”). In these cells, the MSCD plateau level increases throughout prophase I (including after *T*_3_, see the bottom plot of Fig. S7 in Supplementary Information), confirming that the confinement we see beginning at *T*_3_ is specific to homolog pairs.

## Discussion

In this study, we sought to address three major questions: 1) Once “paired”, do two loci remain colocalized? 2) How many interactions are needed to hold homologs in close proximity? 3) How does pairing at one site influence the behavior of nearby sites? Our experimental data, and the theoretical model we developed from it, were able to provide answers to these questions.

We show here that the process of homolog pairing in meiosis is more dynamic than expected from previous observations of static “snapshots”. Our polymer model based on these data revealed that 2-3 linkages per chromosome can act as tethers, confining the otherwise diffusive behavior of distal chromatin, and that these linkages approximate the number of SICs per chromosome. While wild-type strains showed tethered locus mobility at late meiosis, explained in the model by an increased number of linkages, this reduced confinement was not apparent in the *spoii*Δ mutant, suggesting that the linkages confining motion in wild-type represent products of homologous recombination. Finally, these linkages act to restrict the diffusion of adjacent regions of the chromosome, suggesting that they have a functional role in facilitating pairing. These findings highlight how the combination of yeast genetics, *in vivo* single molecule dynamics, and polymer physics modeling can be a powerful tool for understanding complex structural and organizational rearrangements in the nucleus.

### Locus colocalization is dynamic over all stages of meiotic prophase

One of the most surprising results to come from this study was the highly dynamic changes in interlocus distances over time in individual cells and highlights the need to study this dynamic process in live cells. These results build on studies measuring distances between homologous loci using fluorescence *in situ* hybridization (FISH) whereby homologs are paired through multiple-interstitial interactions such that any given locus might be “paired” at any given time, but not all loci are “paired” at once (17, 27). Evidence of dynamic pairing interactions is also seen in meiotic cells of fission yeast (81, 82) and the male *Drosophila* germline (83). By contrast, “somatic” homolog pairing in *Drosophila* showed pairing to be stable such that once paired, the loci remained paired over the course of 8 hours (84). It is likely that features intrinsic to meiotic chromosome architecture and/or telomere-led motion contribute to a more dynamic behavior.

Our biophysical model takes into account predicted contributors to the high level of cell-to-cell heterogeneity that we observed, including the variability in the timing of biological events (e.g. transition from the Rabl configuration, DSB formation and repair), intrinsic cell-to-cell variability in the diffusivity (85), and the formation of linkages that are randomly positioned along homologous chromosomes. Relating single-cell results (characterized in Fig. 5) to ensemble-average behavior (shown in Fig. 7), we show that the *LYS2* and *URA3* loci behave as predicted if located on thermally fluctuating polymers with ~2-4 total linkages in late prophase. This number is far fewer than the combined number of CO and NCOs that have been estimated for these chromosomes (~10-11 and ~7-9) (30). Instead, the number of linkages approximates the number of synapsis initiation complexes (SIC) found along yeast meiotic chromosomes based on the number of Zip2 or Zip3 foci (37, 41). SICs also mark sites in mid prophase that will become Class I COs at late prophase stages (33), or the equivalent of ~ *T*_3_ – *T*_5_ hours in the study. While SIC values have not been empirically determined for chromosomes II and V, we can take numbers reported in these studies and estimate, based on chromosome size, that chromosome II forms ~3-4 SICs and chromosome V forms ~2-3 SICs. The estimated number of SICs for chromosome V is a surprisingly good match for the estimated number of linkages (~3.1-3.36) that best model the behavior of the tagged loci. The number of linkages for chromosome II (~2) is somewhat below expectation. Given the greater length of chromosome II, it is possible that not all SICs (perhaps those more distally located) will serve to limit the confinement radius of tagged loci. Thus, we propose that the establishment of SICs, or at least a subset, representing the sites of future class I COs, will mark the future linkages that limit the extent to which loci diffuse away from one another.

### Local “breathing” of paired chromosome axes may account for dynamic behavior

While the majority of temporal snapshots show only one focus by late meiosis, we also find that the majority of cells are in a “mixed” state. Thus loci that appear “paired” in a snapshot may be moving into and out of proximity. We envision two possible models that could account for this behavior: (1) The sizes of loops attached to chromosome axes undergo dramatic lengthening and shortening, or (2) the two axes “breathe” by transiently moving apart, or some combination of the two. However, these models must account for the large average distance observed between the unpaired homologs, which is about 0.75 μm and continues to reach ~2 μm (the diameter of the nucleus) at late prophase. MSCD data also show that homologs can separate by large fractions of the nucleus in a single time step. What is more, pairing and unpairing is fast, with the dwell time distributions showing that the vast majority of states, both paired and unpaired, are only observed for 30 seconds (the limit of our temporal resolution) before transitioning.

One possibility is that the arrays carried on chromosome loops could be subject to motion by dynamic changes in loop size. Meiotic chromosomes are organized in a loop/axis configuration where DNA is attached to the chromosome axis in a series of loops ranging from 10-50 kb and 28 kb on average (86). Changes in loop size might occur through a loop extrusion mechanism, through dynamic detachment and reattachment of chromatin to the axes (87, 88, 89), or sliding back and forth through ring-shaped cohesin complexes acting as “slip rings” (90). Assuming chromatin compaction in the loops is between 16 nm/kb (77) and 31.6 nm/kb (our model), the ends of the longest extended loops (25 kb) are predicted to be between 400 nm and 800 nm from the chromosome axis. While dynamic changes in loop sizes could contribute in part to the observed behavior, this model is not consistent with the large distances (~200 to 300 kb) between putative linkages or the average distances between foci.

If the SC axes can “breathe” then this would suggest that the SC may be more dynamic than previously anticipated. Several lines of evidence highlight potentially dynamic properties of the SC; first, local separation of the chromosome axes has been observed near recombination complexes in the fungus *Sordaria*, *C. elegans* and *Drosophila* and can be exaggerated in mutant situations (91, 92, 93, 94); second, the transverse filament protein can be exchanged with nuclear pools in yeast and *C. elegans*, particularly near the sites where Class I CO will form (40, 95, 96). The nature of the “breathing” warrants further study since synapsis in *C. elegans* appears to be largely irreversible (97), but increased stability of synapsis may be needed to tether homologs to one another since each chromosome experiences only one crossover (98).

### Locus colocalization is a diffusion-dominated process limited by Spo11-dependent interhomolog tethering

Our biophysical model suggests chromosomes behave as diffusing polymers when pairing is first being established at ~ *T*_4_ hours post transfer to SPM. While telomeres are subject to active motion during meiotic prophase (7), this does not appear to influence interhomolog dynamics where the foci behave as diffusing polymers. A pairing process primarily driven by diffusion would suggest that instead of pushing or pulling chromosome loci together or apart, active telomere motion need only increase fluctuations along the polymer to tacilitate diffusion (i.e. as an effective temperature) (26, 99). Indeed, mutations that abolish telomere-led motion delay but do not eliminate pairing in yeast (4, 26) or in *C. elegans* (99).

If an active mechanism pulled homologous loci together, then we would expect the distributions of pairing times to follow an exponential distribution (100, 101). However, we observed heavy-tailed distributions which suggests that loci are not brought into proximity by means other than polymer diffusion. In addition, it should be noted that we observed very few trajectories that showed hyperdiffusive properties, such as those observed in the case of DSB-dependent directed motion between ALT telomeres (52).

While our model is consistent with foci moving into and out of proximity by diffusion, following the release from the Rabl, and before any linkages (or “tethers”) are established, we cannot rule out that homologous loci interact transiently through reversible biochemical interactions over short time scales (e.g. by multiple rounds of strand invasion (102, 103, 104, 105)). For example, homology could be sensed early on by a “catch-and-release”-like mechanism operating over short time scales, whereby the resected 3’ ends of DSBs undergo multiple invasions before committing to any one of several repair outcomes (43, 106, 107). By our model these transient interactions do not contribute directly to confinement (e.g. behave as “linkages”). Instead, linkages that limit confinement are likely formed later, once the double-Holliday junction intermediate is established (107). It would be interesting to test how tagged loci would behave over both long and short timescales by introducing an inducible DSB nearby as reversible interactions such as this may help to prevent pairing between loci that are only partly homologous (i.e. homeologous) and allow the disentanglement of intertwined chromosomes (108, 109, 110).

### Distal connections alter dynamics at adjacent loci

Our results suggest that once any homolog pair forms a linkage, then that initial connection between the chromosomes will greatly facilitate the interaction of nearby homologous loci. Thus, homolog pairing might happen via a positive feedback mechanism (such as the one proposed in Refs. 16, 47, 48, 72), wherein each random homologous interaction event decreases the colocalization time for all subsequent homologous interactions by forming irreversible bonds via a “ratchet” mechanism that promotes the elongation of the synaptonemal complex as seen in *C. elegans* (99). This appears to be a predominant mechanism for pairing of *Drosophila* chromosomes in somatic cells (84) where the transition from an unpaired to a paired state is a rapid event that occurs in just a few minutes; once “paired” a locus tends to remain paired over long time periods due to homologous “button” interactions (84). This mechanism of pairing is also captured by a thermodynamic phase-transition model proposed in Ref. 15, which incorporates binding molecules to facilitate pairing between homologous sites.

Our experimental results and theoretical model are generally consistent with previous models for homolog pairing (14, 15, 84). However, we note that our observed colocalization dwell-time distributions (Fig. 4a and Supplementary Information, Figs. S3, S4, and S5) exhibit long-time tails that imply the colocalization time is governed by subdiffusive transport associated with polymer dynamics and environmental viscoetasticity. Whereas, specific “button” interactions (84) or binding molecules at pairing centers (14, 15) with significant activation energy would result in colocalization times that are exponentially distributed (i.e. the kinetic model in Fig. 4a), which is inconsistent with our data.

Due to crossover interference, Class I COs are observed less frequently than would be expected from a random distribution (111). Therefore a caveat of our model is that the linkages were randomly distributed across the chromosomes. It will be interesting to add “interfering” crossovers into the model instead to see the effect on pairing. Similarly, it will be interesting to test the effects of tethering in the absence of crossover interference (e.g. in the *zmm* mutants) and other opposing constraints on homolog interactions, including sister chromatid cohesion and telomere-led motion (112).

## Conclusions

We show here that the process of homolog pairing in meiosis is more dynamic than expected from previous observations of static “snapshots” capturing the colocalization of tagged chromosomal loci. We found a large degree of heterogeneous behavior by measuring the mean-squared change in distance of tagged chromosome pairs in individual cells versus ensemble averages. A minimal polymer model reproduces the inter-locus dynamics in premeiotic cells where chromosomes are constrained by the Rabl configuration. The model can also reproduce the physical linkages between homolog pairs that are mediated by the formation and repair of Spo11-induced double strand breaks. These findings highlight how coarse-grained modeling of the basic polymer physics driving chromatin motion can be a powerful tool when dealing with complex structural and organizational rearrangements in the nucleus. With this basic model, we can now begin to add back other variables specific to meiotic chromosomes, such as telomere-led movements, the extension of the synaptonemal complex, crossover interference, and changes in chromosome morphology and compaction over the course of prophase I.

## Materials and methods

### Time course

All yeast strains used were in the SK1 background and are listed in Supplementary Information Fig. S8. Cell synchronization and meiotic induction was performed as described previously (63). Every hour after transfer to sporulation medium, slides were prepared for imaging according to (113), using silicone isolators (Cat. no. JTR20R-2.0, Grace Bio Labs). All of our image processing code is available at https://github.com/ucdavis/SeeSpotRun.

### Imaging

Imaging was performed on a Marianas real time confocal workstation with mSAC + mSwitcher (3i), using a CSU-X1, microlens enhanced, spinning disk unit (Yokogawa). All imaging was performed in a full enclosure environmental chamber preheated to 30 °C, using a microscope incubator (Okolab). Samples were excited with a LaserStack 488nm line (3i), observed using an ALPHA PLAN APO 100X/1.46 OIL objective lens (Zeiss), and photographed using a Cascade QuantEM 512SC camera (Photometrics), with a 0.133μm pixel size. Samples were kept in focus using Definite Focus (Zeiss), capturing up to 41 z-sections (as required to acquire the complete sample thickness), with a 0.25μm step size, every 30s for 50 time points (a total of 25min). Slidebook v5 (3i) was used to run the time-lapse live-cell imaging and export each plane as a separate 16-bit `.tiff` file.

### Video quality control

Videos were excluded from analysis if the quality was so poor as to affect subsequent analysis, with assessments based on signal to noise, signal bleaching, and drift in the z and xy dimensions (Supplementary Information Figs. S15a-c). If drift occurred only at the start or end of the video, and was sufficient to affect image segmentation, then the problematic frames were trimmed from the video. Manual cell segmentation, was performed from a *zt*-MIP (maximum intensity projection, over the z and *t* dimensions) using `distaD_gui.m`, while referring back to the z-MIP video, ignoring overlapping cells and those at the edge of the field of view. Qualitative observations of cell quality were made by referring to the z-MIP video and the position of each cropped cell. Only cells that passed our quality control (Supplementary Information Figs. S15d-j) were included in the subsequent analysis. For inclusion, videos required twice as many live cells as dead (dead/live < 0.5) and > 10 okay cells.

### Spot calling

The position of the fluorescent foci within each cropped cell was detected independently for each time point in the video according to the algorithm described in (114). The raw image intensity data from each cropped cell was filtered with a 3D Gaussian kernel to remove as many noise-related local maxima as possible. Peak localization `(runSpotAnalysistest.m)` was performed through local maxima detection in 3D using image dilation, followed by curvature measurement, which allowed significant peaks to be identified through a cumulative histogram thresholding method. The computational spot calling was manually confirmed in order to remove obvious errors (Supplementary Information Fig. S16) using `conf_gui.m`. If the fitting routine failed to find peaks in more than half the time points for any given cell, that cell was omitted from the analysis.

### Experiment quality control

Experiments with a very poor overall agreement between computational and manual spot calling, with an average difference between detection methods of greater than 10% at each meiotic timepoint, were excluded from analysis. The manual analysis was performed by calling cells as having one or two spots based on a visual assessment of a z-MIP, this was done for three time points from each *T_M_*. Whole experiments were also excluded from the final dataset if the meiotic pairing progression could not be confirmed to exhibit various characteristic properties, such as a single, appropriately timed “nadir”. This was typically due to an experiment lacking sufficient *T_M_* due to exclusion of individual videos.

### Trajectory Analysis

Downstream analysis of the extracted trajectories was performed using a custom Python package (`multi_locus_analysis` (mla) v.0.0.22, see: https://multi-locus-analysis.readthedocs.io/en/latest/). Dwell times were corrected for finite window effects using the method described in (115). Details of the analysis and code used to make plots can be found in the package documentation.

### Analytical Theory

The code used to compute the analytical MSCD curves can also be found in the `wlcsim` codebase under the `wlcsim.analytical.homolog` module (for documentation, see https://wlcsim.readthedocs.io). Briefly, the MSCD calculation is broken down into two cases. In the case where the loci are in between two linkage sites, we treat them as being on an isolated ring polymer whose size is chosen to match the effective ring formed by the two homologous segments holding each locus (which are tethered at either end by the linkage site). This effective ring is outlined in white for “cells” 1 and 4 in Fig. 6. Otherwise, we treat the loci as being on an isolated linear polymer meant to represent the segment of chain running from the end of the first chromosome to one locus, then from that loci to the linkage site, from the linkage site to the other loci, and finally from that loci to the end of the second chromosome. Supplementary Information provides a detailed derivation of the MSCD for these two cases and the value of the plateau MSCD for spherical confined of the polymers.

### Data availability

The raw image data was deposited to the Image Data Resource (http://idr.openmicroscopy.org) under accession number idr0063. The scripts required to reproduce the processed data are available on GitHub (https://github.com/ucdavis/SeeSpotRun); this includes the MATLAB interfaces for spot calling, and the Python scripts for preparing the final xyz position dataset (see Supplementary Dataset 1: finalxyz.csv). The Python module used for downstream analysis also contains the final dataset used in the present study, and can be downloaded from the standard Python repositories by executing `pip install multi_loci_analysis`.

## Supporting information

Supplemental movie "field"

Supplemental movie "mixed"

Supplemental movie "unpaired"

Supplemental movie "paired"

## ACKNOWLEDGMENTS

We thank the lab of Angelika Amon for our FROS strains, and we appreciate helpful discussions and feedback from Amy MacQueen. This work was supported by The National Institutes of Health (NIH), grant: R01GM075119. We thank the Light Microscopy Imaging Facility (Molecular and Cellular Biology, UC, Davis). Financial support for A. J. S. is provided by the National Science Foundation, Physics of Living Systems Program (PHY-1707751). B. B. acknowledges funding support from the NSF Graduate Fellowship program (DGE-1656518) and from an NIH training grant (T32GM008294).

## Notes

The authors declare no conflict of interest.

### Competing Interest Statement

The authors have declared no competing interest.

### Summary of Updates

The manuscript is a revised version of the original "Homologous locus pairing is a transient, diffusion-mediated process in meiotic prophase"

