## Supplemental information for "Diffusion and distal linkages govern interchromosomal dynamics during meiotic prophase"

#### This PDF file includes:

Figs. S1 to S18
Table S1
SI References

### 1. Models for dwell-time distribution

We define three candidate models for the dwell-time distribution based on these models being kinetic, diffusion, and subdiffusion limited. Below, we derive these three models.

**A. Kinetic model.** For dynamics processes that are limited by the rate of a kinetic reaction, the dwell-time distribution is governed by elementary reactions that are individually first-order rate processes. In this picture, times for transitioning between colocalized and separated states exhibit a dwell-time distribution dictated by the set of such reactions. We consider the simplest case where there is either a single elementary reaction or a dominant rate-limiting step among multiple reactions, such that the overall dwell-time distribution is governed by a Poisson distribution. This case coincides with a dwell-time distribution

$$f_t(t) = \frac{1}{t_0} \exp\left(-\frac{t}{t_0}\right), \tag{1}$$

where $t_0$ is a parameter that governs the average time-scale for the transition.

**B. Diffusion model.** To model a diffusion-limited reaction, we adopt a simple picture of a diffusive process in a 1-dimensional landscape of length $L$, symbolizing the diffusive trajectory of the inter-locus distance $x$. This model assumes the particle diffusion occurs in a Newtonian fluid, such that each individual particle exhibits a diffusive mean-square displacement $MSD \sim t^1$. The particle experiences a no-flux boundary at $x = L$ and an absorbing boundary at $x = 0$, which represents the transition from its current state (either separated or colocalized). The particle Green function $G(x|x_0;t)$ gives the probability that if a particle begins at $x_0$ at time zero, it will traverse to $x$ at time $t$. The Green function is governed by the diffusion equation

$$\frac{\partial G(x|x_0;t)}{\partial t} = D\frac{\partial^2 G(x|x_0;t)}{\partial x^2}, \tag{2}$$

with initial condition $G(x|x_0;t=0) = \delta(x-x_0)$ and boundary conditions $G(x=0|x_0;t) = G(x|x_0=0;t) = 0$ and $\partial_x G(x=L|x_0;t) = \partial_x G(x|x_0=L;t) = 0$.

We define the non-dimensional position $\eta = x/L$ and time $\tau = t/t_0$, where $t_0 = L^2/D$, and we identify a set of eigenfunctions

$$\phi_p(\eta) = \sqrt{2}\sin\left[\left(p+\frac{1}{2}\right)\pi\eta\right] \tag{3}$$

for $p = 0, 1, \ldots$. With these definitions, we find the solution to the Green function

$$G(\eta|\eta_0;\tau) = \sum_{p=0}^{\infty} \phi_p(\eta)\phi_p(\eta_0)\exp\left[-\left(p+\frac{1}{2}\right)^2\pi^2\tau\right]. \tag{4}$$

From the Green function, we find the dwell-time distribution for the diffusion model to be

$$f_t(t) = -\frac{1}{t_0}\frac{\partial}{\partial\tau}\left[\int_0^1 d\eta \int_0^1 d\eta_0 G(\eta|\eta_0;\tau)\right] = \frac{2}{t_0}\sum_{p=0}^{\infty}\exp\left[-\left(p+\frac{1}{2}\right)^2\pi^2\tau\right]. \tag{5}$$

**C. Subdiffusion model.** We extend the above model to capture the dwell-time distribution for a transition involving particles that exhibit subdiffusive behavior. Such behavior frequently arises for monomer units in a polymer chain or particles embedded in a viscoelastic environment, which are both invoked in our subsequent detailed models. We assume the particles exhibit a mean-square displacement $MSD \sim t^B$, where $B$ is a power-law exponent between zero and one. Our setup for the subdiffusive model is identical to the diffusive model, and the Green function is governed by a fractional diffusion equation (1, 2)

$$\frac{\partial G(x|x_0;t)}{\partial t} = K D_t^{1-B}\left[\frac{\partial^2}{\partial x^2}G(x|x_0;t)\right] \tag{6}$$

where the fractional-derivative operator is given by

$$D_t^{1-B}W(x,t) = \frac{1}{\Gamma(B)}\frac{\partial}{\partial t}\int_0^t dt' \frac{W(x,t')}{(t-t')^{1-B}}. \tag{7}$$

The subdiffusion coefficient $K$ gives a scale of the rate of subdiffusion.

The subdiffusion model has the same setup and boundary conditions as the diffusion model. In this case, we define the non-dimensional position $\eta = x/L$ and time $\tau = t/t_0$, where $t_0 = (L^2/K)^{1/B}$. With these definitions, we find the solution to the Green function

$$G(\eta|\eta_0;\tau) = \sum_{p=0}^{\infty}\phi_p(\eta)\phi_p(\eta_0)E_{B,1}\left[-\left(p+\frac{1}{2}\right)^2\pi^2\tau^B\right]. \tag{8}$$

where $E_{\alpha,\beta}(x)$ is the Mittag-Leffler function

$$E_{\alpha,\beta}(x) = \sum_{j=0}^{\infty}\frac{x^j}{\Gamma(\beta+\alpha j)}. \tag{9}$$

From the Green function, we find the dwell-time distribution for the diffusion model to be

$$f_t(t) = -\frac{1}{t_0}\frac{\partial}{\partial \tau}\left[\int_0^1 d\eta \int_0^1 d\eta_0 G(\eta|\eta_0;\tau)\right] = \frac{2}{t_0}\tau^{B-1}\sum_{p=0}^{\infty} E_{B,B}\left[-\left(p+\frac{1}{2}\right)^2 \pi^2 \tau^B\right]. \quad [10]$$

We note that the subdiffusion model reduces to the diffusion model in the limit $B \to 1$, as this limit coincides with diffusion in a Newtonian fluid.

### 2. Theoretical model of linked viscoelastic Rouse polymers

We consider a flexible polymer chain that is subjected to Brownian forces (3). Our goals is to determine specific dynamic properties of a linear and ring Gaussian chain. For example, we consider the dynamic motion of two chain segments relative to each other. We define this relative motion as the mean-square change in distance (MSCD). In this development, we present a detailed derivation for the linear chain, and we provide the results for the ring polymer.

We start by defining a discrete polymer chain with bead positions $\vec{r}^{(n)}$, where $n$ runs from 0 to $n_\mathrm{b} - 1$. Each bead is connected to their neighboring beads by Hookean springs, resulting in a potential force on the $n$-th bead $\vec{f}_E^{(n)}$ that is given by

$$\vec{f}_E^{(n)} = \begin{cases} \frac{3k_BT}{gb^2}\left(\vec{r}^{(n+1)} - 2\vec{r}^{(n)} + \vec{r}^{(n-1)}\right) & n = 1,\ldots,n_\mathrm{b}-2 \\ \frac{3k_BT}{gb^2}\left(\vec{r}^{(1)} - \vec{r}^{(0)}\right) & n = 0 \\ -\frac{3k_BT}{gb^2}\left(\vec{r}^{(n_\mathrm{b}-1)} - \vec{r}^{(n_\mathrm{b}-2)}\right) & n = n_\mathrm{b}-1 \end{cases} \quad [11]$$

where $b$ is the Kuhn statistical segment length of the polymer (3) and $g$ is the number of Kuhn lengths per bead. We set $g = N/n_\mathrm{b}$, giving a total chain length of $N$ Kuhn lengths.

The fractional Langevin equation of motion for the $n$th bead is given by

$$g\xi \int_0^t dt' K(t-t')\frac{d\vec{r}^{(n)}(t')}{dt} = \vec{f}_E^{(n)} + \vec{f}_B^{(n)} \quad [12]$$

where the memory kernel is given by

$$K(t-t') = \frac{(2-\alpha)(1-\alpha)}{|t-t'|^{\alpha}}. \quad [13]$$

The drag coefficient $\xi$ is defined as the viscoelastic drag per Kuhn length, and the power-law scaling $\alpha$ characterizes the relative impact of viscous and elastic forces from the environment ($\alpha = 1$ is purely viscous and $\alpha = 0$ is purely elastic). We now take the limit of $n_\mathrm{b} \to \infty$ for fixed chain length $N$, resulting in a continuous-chain representation of the chain $\vec{r}(n,t)$ where $n$ is a path-length variable that runs from 0 to $N$. The Langevin equation for the chain is given by

$$\xi \int_0^t dt' K(t-t')\frac{d\vec{r}^{(n)}(n,t')}{dt} = \frac{3k_BT}{b^2}\frac{\partial^2\vec{r}(n,t)}{\partial n^2} + \vec{f}_B(n,t), \quad [14]$$

which is subjected to the end conditions

$$\frac{\partial \vec{r}(n=0,t)}{\partial n} = \frac{\partial \vec{r}(n=N,t)}{\partial n} = 0. \quad [15]$$

The Brownian forces satisfy the fluctuation dissipation theorem

$$\langle \vec{f}_B(n,t)\vec{f}_B(n',t')\rangle = 2k_BT\xi\delta(n-n')K(t-t')\mathbf{I}. \quad [16]$$

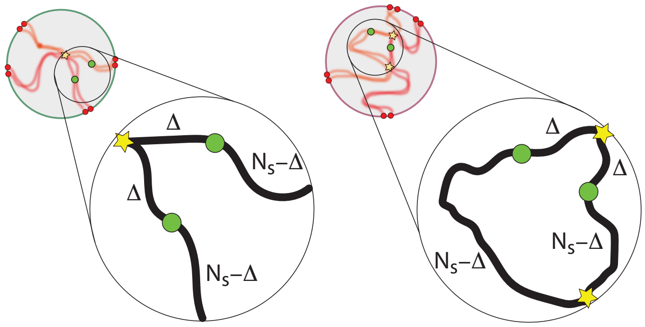


**Fig. S1.** Schematic representation of the linear and ring Rouse model with definition of segments for MSCD calculation. The effective polymer representation on the left captures instances where the visualized locus (*e.g. URA3* on chr. V) has a linkage on one side but no linkage on the other, resulting in an effectively linear chain. The effective polymer representation on the right captures instances where the visualized locus (*e.g. URA3* on chr. V) has a linkage on both sides, making it part of an interior ring. The total chain length is $N = 2N_s$, and we consider two points on the chain located at $n_1 = N_s - \Delta = N/2 - \Delta$ and $n_2 = N_s + \Delta = N/2 + \Delta$.

It is convenient to define a set of normal coordinates that effectively decouple the interactions implicit within the equation of motion (Eq. 14). We define the normal modes

$$\phi_p(n) = \begin{cases} \sqrt{2}\cos\left(\frac{\pi p n}{N}\right), & p = 1, 2, \ldots \\ 1, & p = 0. \end{cases} \quad [17]$$

These modes represent a complete basis set that satisfy the boundary conditions for $\vec{r}(n,t)$. Orthogonality is demonstrated by noting

$$\int_0^N dn \phi_p(n)\phi_{p'}(n) = N\delta_{p,p'}. \quad [18]$$

The amplitude of the $p$-th mode $\vec{X}_p(t)$ is given by

$$\vec{X}_p(t) = \frac{1}{N}\int_0^N dn\, \vec{r}(n,t)\phi_p(n), \quad [19]$$

and the inversion back to chain coordinates is written as

$$\vec{r}(n,t) = \sum_{p=0}^{\infty} \vec{X}_p(t)\phi_p(n). \quad [20]$$

Upon performing a transform to normal coordinates, we find the governing equation of motion

$$\xi N \int_0^t dt' K(t-t') \frac{d\vec{X}_p(t')}{dt} = -k_p \vec{X}_p + \vec{f}_{B_p} \quad [21]$$

where $k_p = \frac{3\pi^2 k_B T}{b^2 N} p^2$. The $p$-mode Brownian force $\vec{f}_{B_p}$ is given by

$$\vec{f}_{B_p} = \int_0^N dn\, \vec{f}_B(n,t)\phi_p(n) \quad [22]$$

and satisfies the fluctuation dissipation theorem

$$\langle \vec{f}_{B_p}(t)\vec{f}_{B_{p'}}(t')\rangle = 2k_B T \xi N \delta_{pp'} K(t-t')\mathbf{I}. \quad [23]$$

A similar derivation for a ring polymer results in a treatment that is identical to the linear chain, but the normal modes for the ring polymer are continuous across the ends [i.e. $\vec{r}(n=0,t) = \vec{r}(n=N,t)$]. Specifically, the complete normal-mode set is separated into even and odd functions, respectively defined by

$$\phi_p^{(e)}(n) = \begin{cases} \sqrt{2}\cos\left(\frac{2p\pi n}{N}\right), & p = 1, 2, \ldots \\ 1, & p = 0. \end{cases} \quad [24]$$

and

$$\phi_p^{(o)}(n) = \sqrt{2}\sin\left(\frac{2\pi p n}{N}\right), \quad p = 1, 2, \ldots \quad [25]$$

The even and odd normal modes satisfy the equation of motion defined in Eq. 21 with $k_p = \frac{12\pi^2 k_B T}{b^2 N} p^2$

**A. Mean-square displacement (MSD) for linear polymers.** The mean-square displacement (MSD) of a segment of the polymer chain is define a

$$\text{MSD} = \langle (\vec{r}(n,t) - \vec{r}(n,0))^2 \rangle \quad [26]$$

for the $n$th segment of the chain with total length $N$. We insert our normal-mode representation into Eq. 26, resulting in the expression

$$\text{MSD} = \sum_{p=0}^{\infty}\sum_{p'=0}^{\infty} \langle \left(\vec{X}_p(t) - \vec{X}_p(0)\right) \cdot \left(\vec{X}_{p'}(t) - \vec{X}_{p'}(0)\right)\rangle \phi_p(n)\phi_{p'}(n) \quad [27]$$

The equation of motion (Eq. 21) can be used to determine the correlation function $\langle \vec{X}_p(t) \cdot \vec{X}_{p'}(0)\rangle$ (detailed discussion is found in Ref. (4)). This results in the expression

$$\langle \vec{X}_p(t) \cdot \vec{X}_{p'}(0)\rangle = 3\frac{k_B T}{k_p} E_{\alpha,1}\left[-\frac{k_p}{N\xi\Gamma(3-\alpha)} t^{\alpha}\right] \delta_{pp'} \quad [28]$$

for $p \geq 1$, where $E_{\alpha,\beta}(x)$ is the Mittag-Leffler function

$$E_{\alpha,\beta}(x) = \sum_{j=0}^{\infty} \frac{x^j}{\Gamma(\beta + \alpha j)}. \quad [29]$$

For $p = 0$, we find the expression

$$\langle (\vec{X}_0(t) - \vec{X}_0(0))^2 \rangle = \frac{3k_BT}{N\xi} \frac{\sin(\alpha\pi)}{\pi(1-\alpha/2)(1-\alpha)\alpha} t^{\alpha}. \quad [30]$$

We focus on the MSD for the midpoint of a linear chain, thus $n = N/2$. Inserting this into our definition of MSD results in the expression for the MSD of the midpoint of a linear chain

$$\begin{aligned} \text{MSD} &= 6\frac{k_BT}{\xi N} t + \sum_{p\,\text{even}} 12\frac{k_BT}{k_p}\left\{1 - E_{\alpha,1}\left[-\frac{k_p}{N\xi\Gamma(3-\alpha)}t^{\alpha}\right]\right\} && [31] \\ &= 6\frac{k_BT}{\xi N} t + \sum_{p=1}^{\infty} 12\frac{k_BT}{k_{2p}}\left\{1 - E_{\alpha,1}\left[-\frac{k_{2p}}{N\xi\Gamma(3-\alpha)}t^{\alpha}\right]\right\} && [32] \end{aligned}$$

**B. Mean-squared change in distance (MSCD) for linear and ring polymers.** We now consider the mean-square change in distance (MSCD) for a linear polymer chain. This quantity is defined as

$$\text{MSCD} = \langle \left(\Delta\vec{R}(t) - \Delta\vec{R}(0)\right)^2 \rangle \quad [33]$$

where $\Delta\vec{R}(t) = \vec{r}(N/2+\Delta, t) - \vec{r}(N/2-\Delta, t)$ where the total chain length is $N = 2N_s$. We insert our normal-mode representation into Eq. 33 to find

$$\text{MSCD} = \sum_{p=1}^{\infty}\sum_{p'=1}^{\infty} \langle \left(\vec{X}_p(t) - \vec{X}_p(0)\right) \cdot \left(\vec{X}_{p'}(t) - \vec{X}_{p'}(0)\right) \rangle \left[\phi_p(N/2+\Delta) - \phi_p(N/2-\Delta)\right]\left[\phi_{p'}(N/2+\Delta) - \phi_{p'}(N/2-\Delta)\right] \quad [34]$$

The equation of motion (Eq. 21) can be used to determine the correlation function $\langle \vec{X}_p(t) \cdot \vec{X}_{p'}(0) \rangle$ (detailed discussion is found in Ref. (4)). This results in the expression

$$\langle \vec{X}_p(t) \cdot \vec{X}_{p'}(0) \rangle = 3\frac{k_BT}{k_p} E_{\alpha,1}\left[-\frac{k_p}{N\xi\Gamma(3-\alpha)}t^{\alpha}\right]\delta_{pp'} \quad [35]$$

Inserting this into our definition of MSCD results in the expression for the linear chain

$$\begin{aligned} \text{MSCD}^{(\text{linear})} &= \sum_{p\,\text{odd}} 48\frac{k_BT}{k_p}\left\{1 - E_{\alpha,1}\left[-\frac{k_p}{N\xi\Gamma(3-\alpha)}t^{\alpha}\right]\right\}\sin^2\left(\frac{\pi p\Delta}{N}\right) && [36] \\ &= \sum_{p=0}^{\infty} 48\frac{k_BT}{k_{2p+1}}\left\{1 - E_{\alpha,1}\left[-\frac{k_{2p+1}}{N\xi\Gamma(3-\alpha)}t^{\alpha}\right]\right\}\sin^2\left[\frac{\pi(2p+1)\Delta}{N}\right] && [37] \end{aligned}$$

where $k_p = \frac{3\pi^2 k_BT}{b^2 N}p^2$ for the linear chain.

We follow a parallel derivation for the ring polymer. We note that only the odd set of normal modes contribute to MSCD for the ring polymer. Taking similar steps as in the linear case, we arrive at the expression for the ring polymer

$$\text{MSCD}^{(\text{ring})} = \sum_{p=1}^{\infty} 48\frac{k_BT}{k_p}\left\{1 - E_{\alpha,1}\left[-\frac{k_p}{N\xi\Gamma(3-\alpha)}t^{\alpha}\right]\right\}\sin^2\left(\frac{2\pi p\Delta}{N}\right) \quad [38]$$

where $k_p = \frac{12\pi^2 k_BT}{b^2 N}p^2$ for the ring polymer.

The code used to compute the analytical MSCD curves can also be found in the `wlcsim` codebase under the `wlcsim.analytical.homolog` module (for documentation, see https://wlcsim.readthedocs.io).

**C. MSCD plateau due to nuclear confinement.** We consider a flexible Gaussian chain in a spherical confinement of radius $a$. Within the confinement, the potential is set to zero, i.e. $V = 0$ for $R < a$. The Green function is zero outside the confinement, so we enforce the confinement by setting $G = 0$ at $R = a$ and $R_0 = a$. The Green function for the 3-D Gaussian Chain within the confinement obeys

$$\frac{\partial G(\vec{R}, N|\vec{R}_0, 0)}{\partial N} = \frac{b^2}{6}\vec{\nabla}^2 G(\vec{R}, N|\vec{R}_0, 0) \quad [39]$$

with the boundary condition

$$\begin{aligned} G(\vec{R} = a \cdot \hat{e}_R, N|\vec{R}_0, 0) = 0 \\ G(\vec{R}, N|\vec{R}_0 = a \cdot \hat{e}_{R_0}, 0) = 0 \end{aligned} \quad [40]$$

where $\hat{e}_R$ and $\hat{e}_{R_0}$ are unit vectors in the direction of $\vec{R}$ and $\vec{R}_0$, respectively.

We use separation of variables to solve for the Green function within the confinement. We define the position-dependent wavefunction $\psi_{nlm} = R_{nl}(R)Y_l^m(\hat{e})$. We non-dimensionalize the radial distance by $a$, defining the dimensionless position vector $\vec{r} = \vec{R}/a$. With this definition, we determine the Green function by solving the appropriate eigenfunction problem, given by

$$\mathcal{H}\psi_{nlm} = \vec{\nabla}^2\psi_{nlm} = -\lambda_{nlm}^2\psi_{nlm} \quad [41]$$

Inserting our wavefunction into this equation gives the governing equation

$$\left[\frac{1}{r^2}\frac{d}{dr}r^2\frac{d}{dr} - \frac{l(l+1)}{r^2}\right]R_{nl}(r) = -\lambda_{nl}^2 R_{nl}(r) \quad [42]$$

where we note that $\lambda$ only depends on the indices $n$ and $l$. This adopts the form of the spherical Bessel differential equation, leading to solutions of the form $R_{nl} = Aj_l(\lambda_{nl}r) + Bn_l(\lambda_{nl}r)$, where $j_l$ and $y_l$ are the spherical Bessel functions of the first and second kind, respectively. Since $r = 0$ is part of the domain of interest, we exclude $y_l$ from our solution, since these functions diverge as $r \to 0$. The boundary condition is satisfied by forcing $R(r = 1) = 0$, leading to the condition $j_l(\lambda_{nlm}) = 0$.

We then write the full solution for the Green function in the form

$$G(\vec{R}|\vec{R}_0; N) = \sum_{nlm}\psi_{nlm}(\vec{R})\psi_{nlm}^*(\vec{R}_0)\exp\left[-\frac{1}{6}\lambda_{nl}^2\frac{b^2N}{a^2}\right] \quad [43]$$

where

$$\psi_{nlm}(\vec{R}) = A_{nlm}j_l(\lambda_{nl}R/a)Y_l^m(\hat{e}) \quad [44]$$

The normalization constant $A_{nlm}$ dictates that

$$\int d\hat{e}\int_0^a dRR^2\psi_{nlm}(\vec{R})\psi_{n'l'm'}^*(\vec{R}) = \delta_{nn'}\delta_{ll'}\delta_{mm'} \quad [45]$$

***C.1. Chain segmental density and average position for free ends.*** From this Green function, we now determine the segmental density and average position of a polymer chain within a confinement. Specifically, we address the behavior for both a chain with free ends and a chain with both ends attached to the sphere surface (as applied to organization within the yeast nucleus during prophase I of meiosis). We define the segmental density for a segment at position $\Delta$ for a chain of total length $N$. For free ends, the segment density is given by

$$\rho_{\text{free}}(\vec{R}_\Delta) = \frac{1}{\mathcal{N}_{\text{free}}}\int d\vec{R}d\vec{R}_0 G(\vec{R}|\vec{R}_\Delta; N-\Delta)G(\vec{R}_\Delta|\vec{R}_0;\Delta) \quad [46]$$

$$= \frac{4a^{-6}}{\mathcal{N}_{\text{free}}}\int_0^a dRR^2\int_0^a dR_0R_0^2\sum_{n=1}^{\infty}\sum_{n_0=1}^{\infty}\frac{\sin(n\pi R/a)}{R}\frac{\sin(n\pi R_\Delta/a)}{R_\Delta}\frac{\sin(n_0\pi R_\Delta/a)}{R_\Delta}\frac{\sin(n_0\pi R_0/a)}{R_0} \quad [47]$$

$$\times\exp\left[-\frac{1}{6}(n\pi)^2\frac{b^2(N-\Delta)}{a^2}\right]\exp\left[-\frac{1}{6}(n_0\pi)^2\frac{b^2\Delta}{a^2}\right] \quad [48]$$

$$= \frac{4}{\mathcal{N}_{\text{free}}}\sum_{n=1}^{\infty}\sum_{n_0=1}^{\infty}\frac{(-1)^n}{(n\pi)}\frac{(-1)^{n_0}}{(n_0\pi)}\frac{\sin(n\pi R_\Delta/a)\sin(n_0\pi R_\Delta/a)}{R_\Delta^2} \quad [49]$$

$$\times\exp\left[-\frac{1}{6}(n\pi)^2\frac{b^2(N-\Delta)}{a^2}\right]\exp\left[-\frac{1}{6}(n_0\pi)^2\frac{b^2\Delta}{a^2}\right] \quad [50]$$

where $\mathcal{N}_{\text{free}}$ is a normalization constant that ensures $\int d\vec{R}_\Delta\rho_{\text{free}}(\vec{R}_\Delta) = 1$. With this, we find the normalization constant to be

$$\mathcal{N}_{\text{free}} = 8\pi a^3\sum_{n=1}^{\infty}\frac{1}{(n\pi)^2}\exp\left[-\frac{1}{6}(n\pi)^2\frac{b^2N}{a^2}\right] \quad [51]$$

From this, we find the average squared position of the segment within the sphere to be

$$\langle R_\Delta^2\rangle_{\text{free}} = \frac{16\pi a^5}{\mathcal{N}_{\text{free}}}\sum_{n=1}^{\infty}\sum_{n_0=1}^{\infty}\frac{(-1)^n}{(n\pi)}\frac{(-1)^{n_0}}{(n_0\pi)}I_{nn_0}\exp\left[-\frac{1}{6}(n\pi)^2\frac{b^2(N-\Delta)}{a^2}\right]\exp\left[-\frac{1}{6}(n_0\pi)^2\frac{b^2\Delta}{a^2}\right] \quad [52]$$

where the integral factor $I_{nn_0}$ is given by

$$I_{nn_0} = \frac{4nn_0(-1)^n(-1)^{n_0}}{(\pi n^2 - \pi n_0^2)^2} \quad [53]$$

if $n \neq n_0$, and

$$I_{nn} = \frac{1}{6} - \frac{1}{4\pi^2n^2} \quad [54]$$

for the case $n = n_0$.

***C.2. Chain segmental density and average position for attached ends.*** We now find the segmental density for the case where the chain ends are attached to the sphere surface. We assume the ends are free to move on the sphere surface, such that $\vec{R} = (a-\epsilon)\hat{e}$ and $\vec{R}_0 = (a-\epsilon)\hat{e}_0$. Since the Green function approaches zero on the sphere surface, we set the position a small distance $\epsilon$ within the sphere. We will take the limit $\epsilon \to 0$ after we find the average to determine the limiting behavior for surface attachment.

With this development, we find the segmental density within the sphere to be

$$\rho_{\text{surf}}(\vec{R}_\Delta) = \frac{1}{\mathcal{N}_{\text{surf}}} \int d\hat{e} d\hat{e}_0 G(\vec{R} = (a-\epsilon)\hat{e}|\vec{R}_\Delta; N-\Delta) G(\vec{R}_\Delta|\vec{R}_0 = (a-\epsilon)\hat{e}_0; \Delta) \quad [55]$$

$$= \frac{4a^{-6}}{\mathcal{N}_{\text{surf}}} \sum_{n=1}^{\infty} \sum_{n_0=1}^{\infty} \frac{\sin[n\pi(a-\epsilon)/a]}{(a-\epsilon)} \frac{\sin(n\pi R_\Delta/a)}{R_\Delta} \frac{\sin(n_0\pi R_\Delta/a)}{R_\Delta} \frac{\sin[n_0\pi(a-\epsilon)/a]}{(a-\epsilon)} \quad [56]$$

$$\times \exp\left[-\frac{1}{6}(n\pi)^2 \frac{b^2(N-\Delta)}{a^2}\right] \exp\left[-\frac{1}{6}(n_0\pi)^2 \frac{b^2\Delta}{a^2}\right] \quad [57]$$

$$= \frac{4\epsilon^2 a^{-10}}{\mathcal{N}_{\text{surf}}} \sum_{n=1}^{\infty} \sum_{n_0=1}^{\infty} (-1)^n (n\pi)(-1)^{n_0}(n_0\pi) \frac{\sin(n\pi R_\Delta/a)\sin(n_0\pi R_\Delta/a)}{R_\Delta^2} \quad [58]$$

$$\times \exp\left[-\frac{1}{6}(n\pi)^2 \frac{b^2(N-\Delta)}{a^2}\right] \exp\left[-\frac{1}{6}(n_0\pi)^2 \frac{b^2\Delta}{a^2}\right] \quad [59]$$

We find the normalization constant to be

$$\mathcal{N}_{\text{surf}} = 8\pi\epsilon^2 a^{-7} \sum_{n=1}^{\infty} (n\pi)^2 \exp\left[-\frac{1}{6}(n\pi)^2 \frac{b^2 N}{a^2}\right] \quad [60]$$

From this, we find the average squared position of the segment within the sphere to be

$$\langle R_\Delta^2 \rangle_{\text{surf}} = \frac{16\pi\epsilon^2 a^{-5}}{\mathcal{N}_{\text{surf}}} \sum_{n=1}^{\infty} \sum_{n_0=1}^{\infty} (-1)^n (n\pi)(-1)^{n_0}(n_0\pi) I_{nn_0} \exp\left[-\frac{1}{6}(n\pi)^2 \frac{b^2(N-\Delta)}{a^2}\right] \exp\left[-\frac{1}{6}(n_0\pi)^2 \frac{b^2\Delta}{a^2}\right] \quad [61]$$

where the integral factor $I_{nn_0}$ is given by

$$I_{nn_0} = \frac{4nn_0(-1)^n(-1)^{n_0}}{(\pi n^2 - \pi n_0^2)^2} \quad [62]$$

if $n \neq n_0$, and

$$I_{nn} = \frac{1}{6} - \frac{1}{4\pi^2 n^2} \quad [63]$$

for the case $n = n_0$.

### 3. Supplemental data and analysis

**Table S1. Genomic properties of the LYS2 and URA3 loci**

| | LYS2 | URA3 |
|---|---|---|
| Chromosome | II | V |
| Total chromosome length | 813 kb | 577 kb |
| Approximate position of tag | ∼470 kb | ∼116 kb |
| Centromere position | 238 kb | 152 kb |
| Distance from centromere | ∼232 kb | ∼36 kb |
| Estimated distance from centromere (No. of Kuhn lengths) | 29.6 | 4.5 |
| Distance from nearest telomere | ∼343 kb | ∼116 kb |
| Estimated distance from closest telomere (No. of Kuhn lengths) | 43.3 | 14.8 |
| Average No. of CO/NCO | ∼6-7/∼4 | ∼4-5/∼3 |
| Estimated No. of SIC (Class I CO) | 3-4 | 2-3 |
| Estimated distance between linkages | ∼203-271 kb | 192-288 kb |

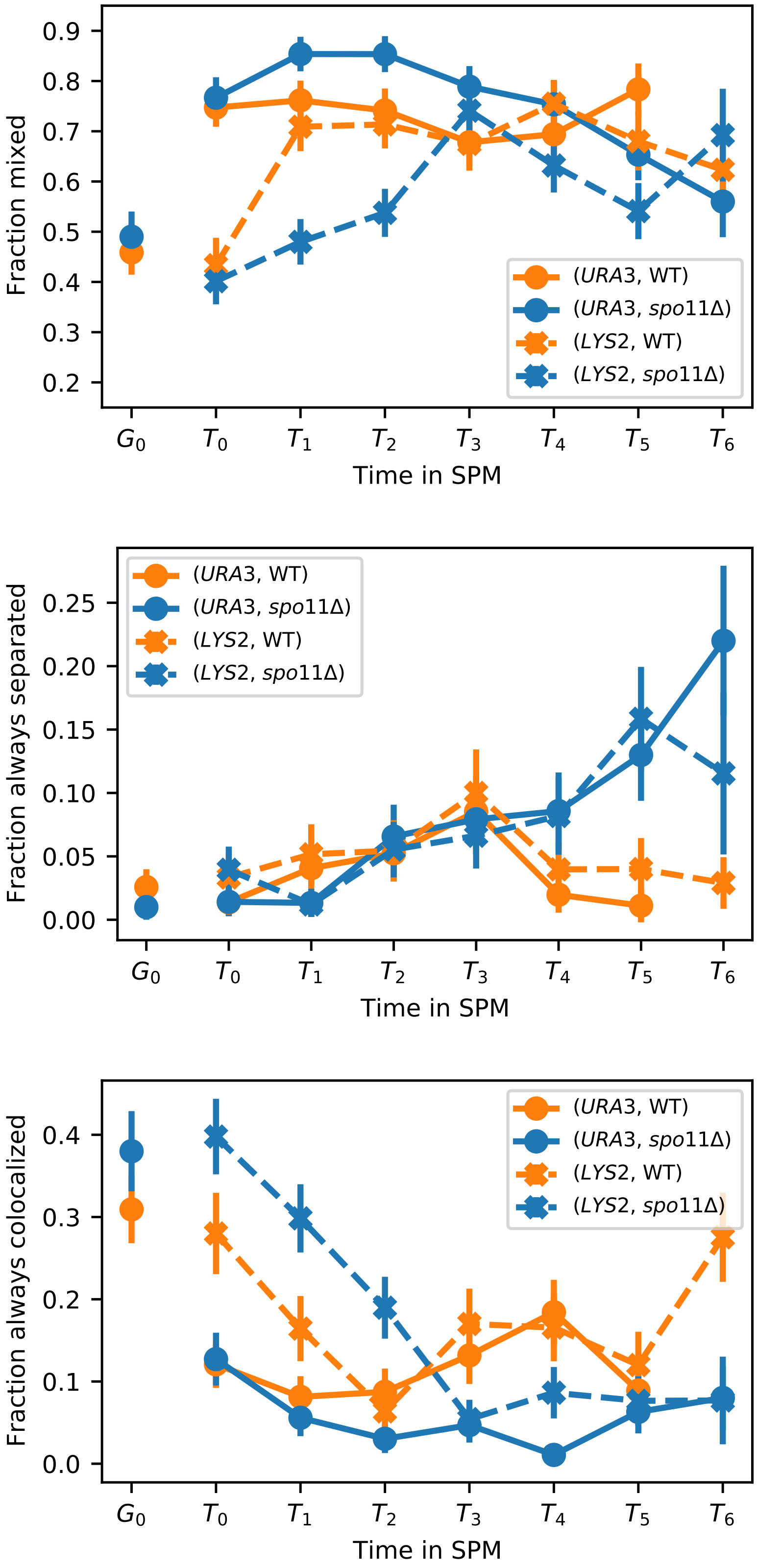


**Fig. S2.** Comparison between fractions of (a) frames where loci are observed to be colocalized, (b) number of cells where both colocalized and separated states are observed, and (c) the fraction of cells that are always separated.

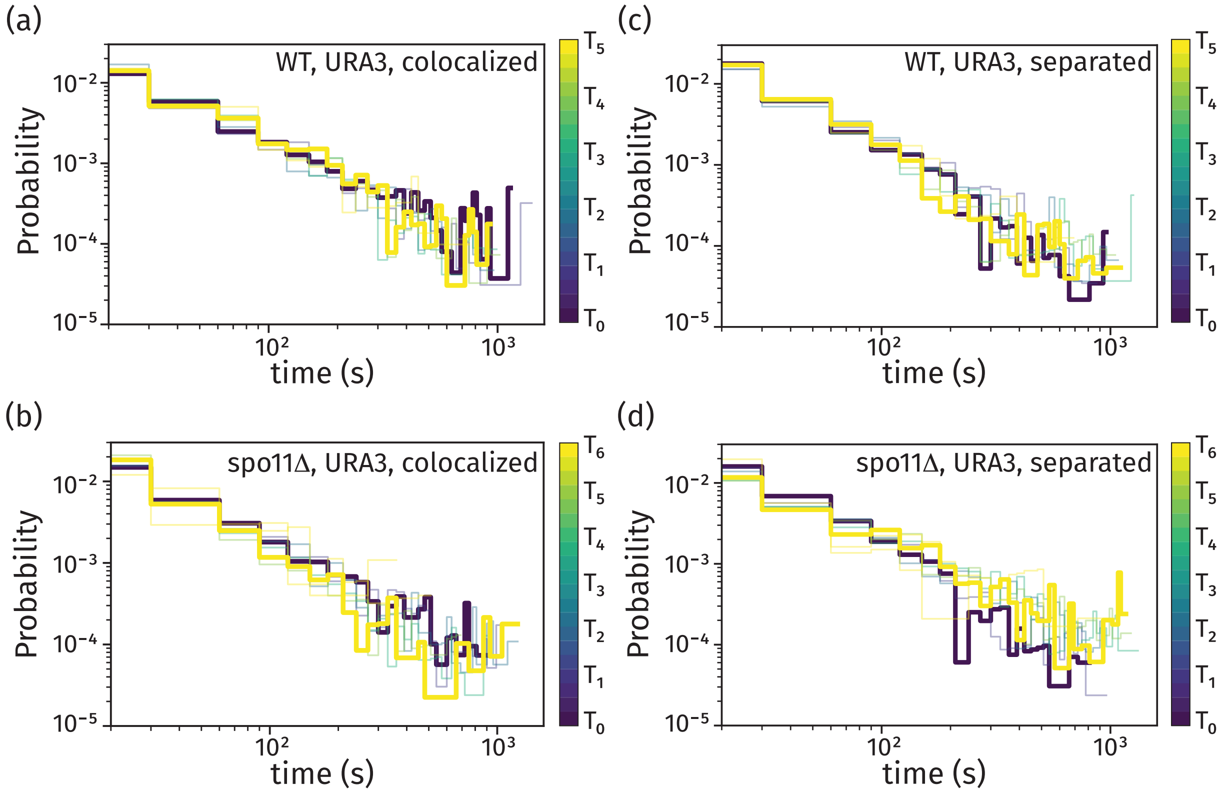


**Fig. S3.** Histograms of dwell times in the colocalized and separated states for the *URA3* locus. One histogram per stage in meiosis is shown, colored by the time since transfer to sporulation media. Experimental data is shown for the colocalized state, including data for wild-type (a) and *spo11*Δ (b) strains, and separated state for wild-type (c) and *spo11*Δ (d) strains.

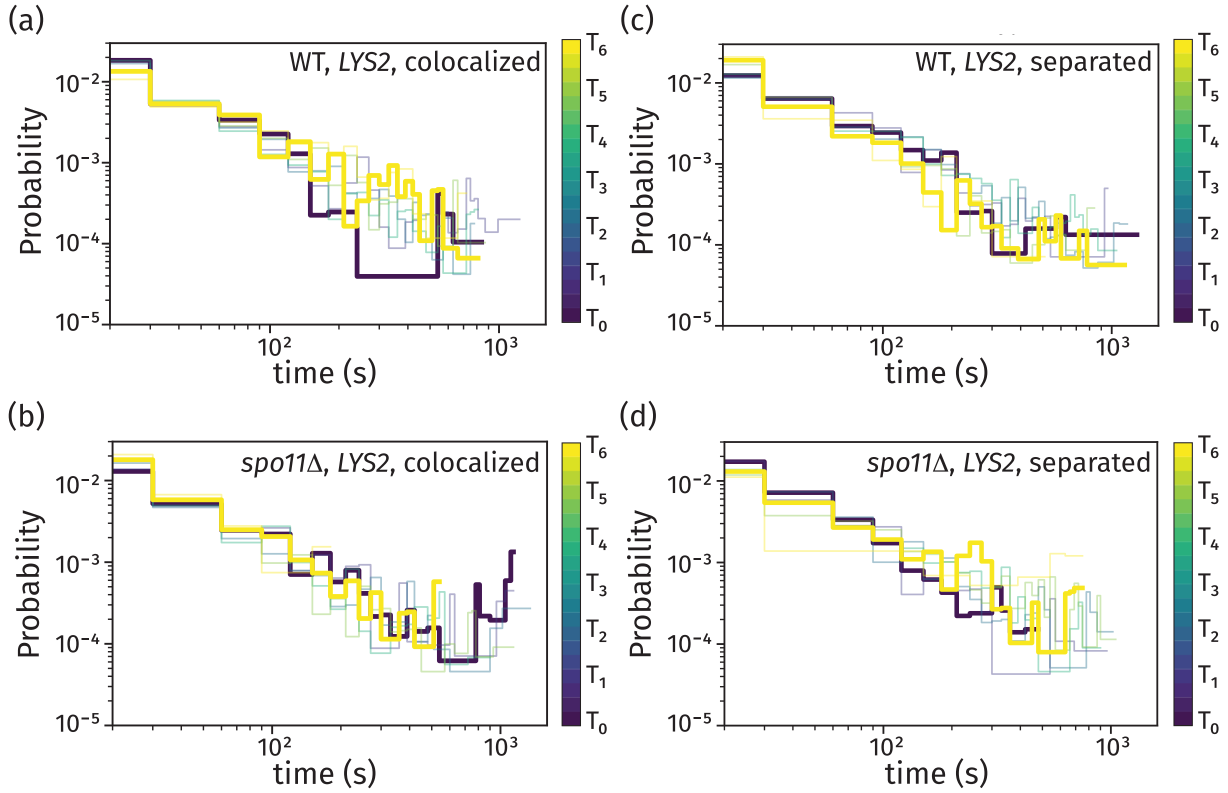


**Fig. S4.** Histograms of dwell times in the colocalized and separated states for the *LYS2* locus. One histogram per stage in meiosis is shown, colored by the time since transfer to sporulation media. Experimental data is shown for the colocalized state, including data for wild-type (a) and *spo11*Δ (b) strains, and separated state for wild-type (c) and *spo11*Δ (d) strains.

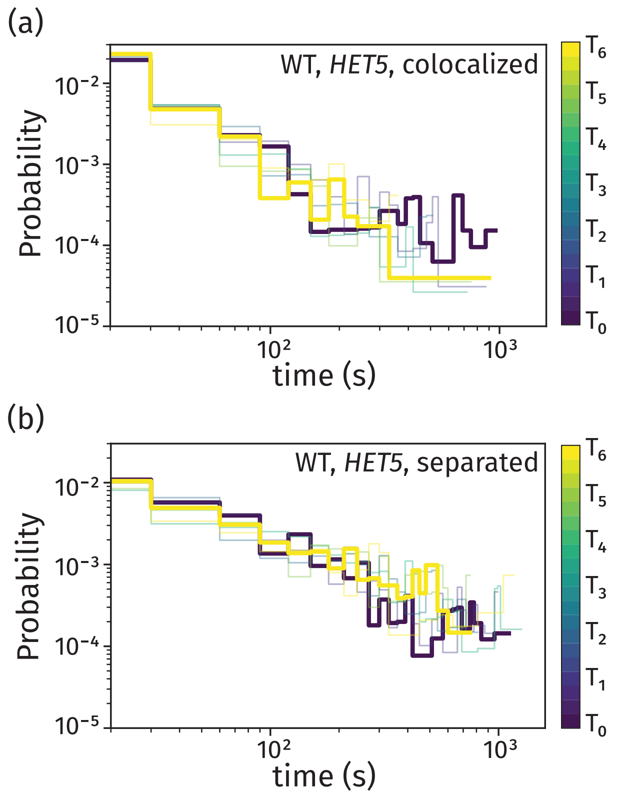


**Fig. S5.** Histograms of dwell times in the colocalized (a) and separated (b) states for the *HET5* locus. One histogram per stage in meiosis is shown, colored by the time since transfer to sporulation media.

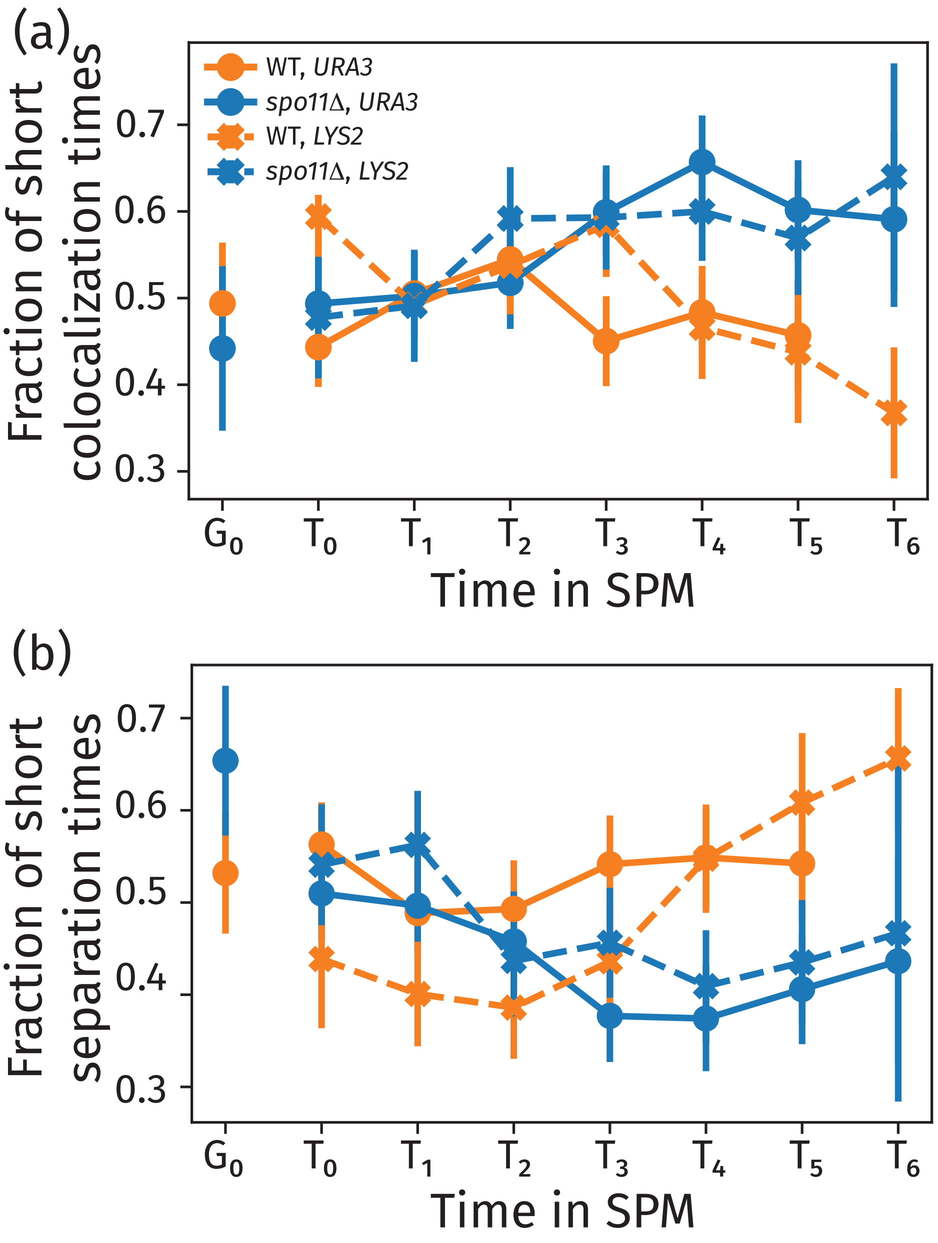


**Fig. S6.** We show the height of the first bar of the waiting time histogram for each strain. The height of the first bar of the colocalization time distribution roughly corresponds to what fraction of the time the interactions are fast (top). Similarly, the height of the first bar of the separation time distribution corresponds to the fraction of the separation times that are fast. We see that in WT, less transient colocalization events occur and separation times are faster as meiosis progresses, but that is no longer true for the *spo11*Δ mutant.

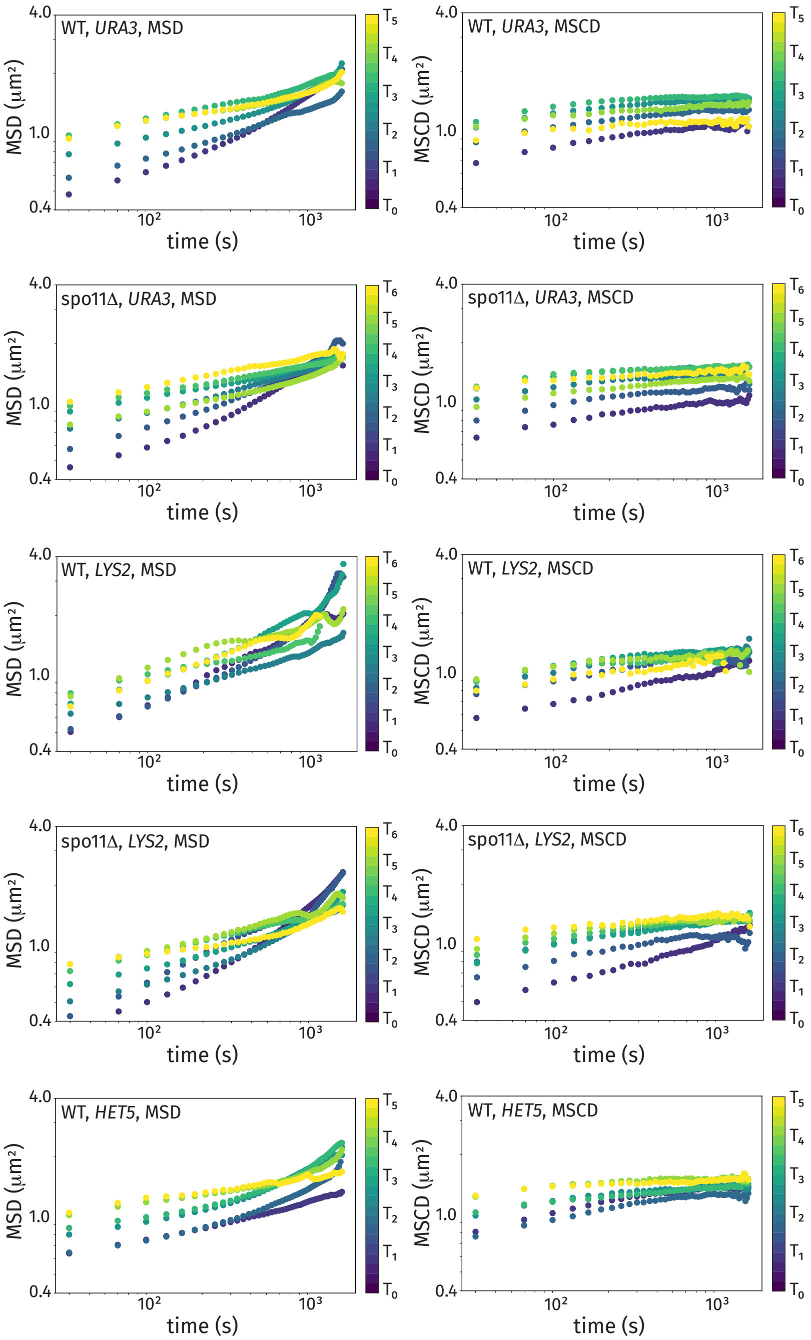


**Fig. S7.** Comparison between the mean-square change in distance (MSCD) and mean-square displacment (MSD) for both the *URA3* and *LYS2* loci in wild-type cells.

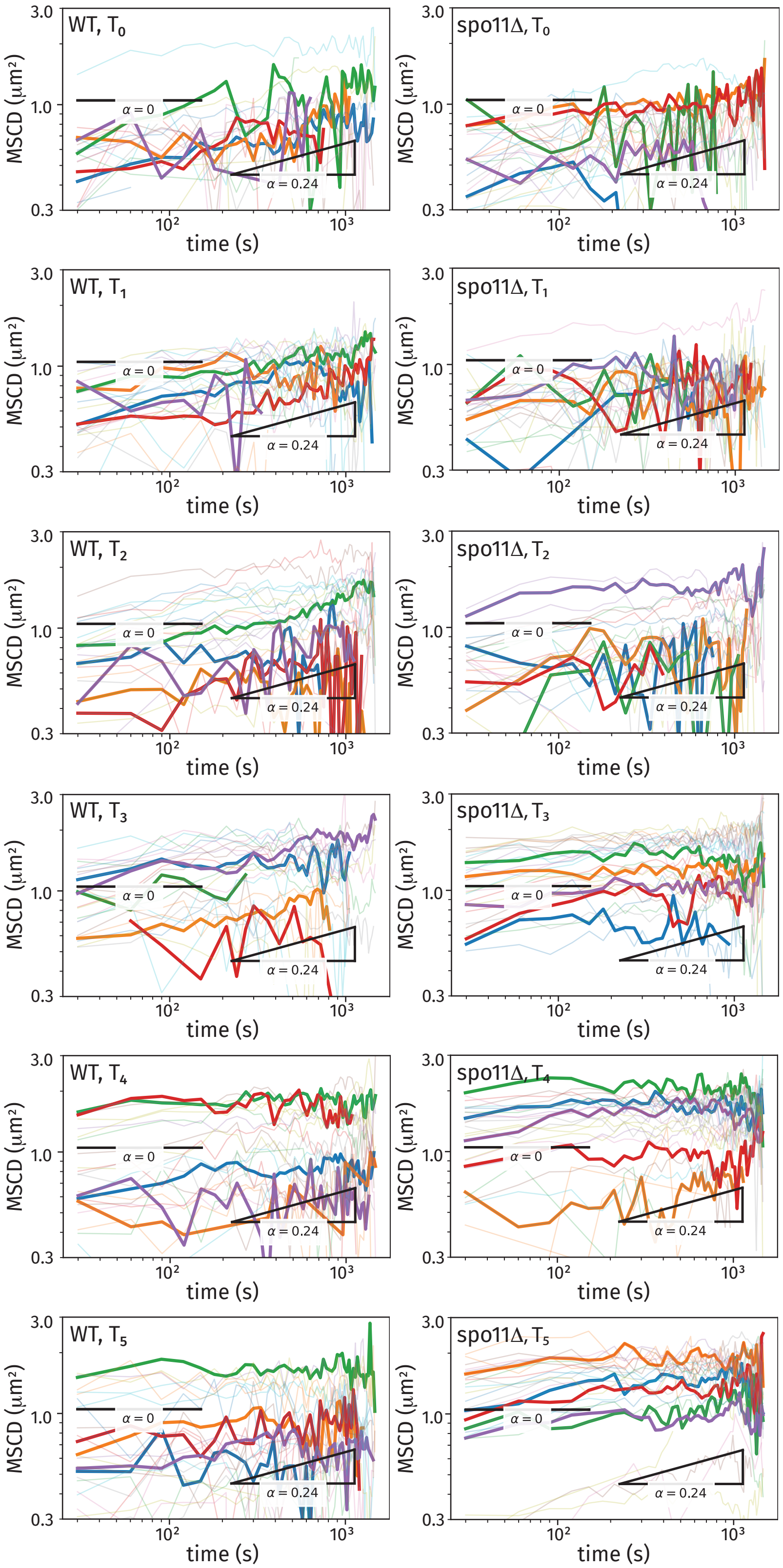


**Fig. S8.** Single-cell MSCD plots for the *URA3* locus for chronological stage $T_0$ to $T_5$ (same presentation as in Fig. 5 of the manuscript).

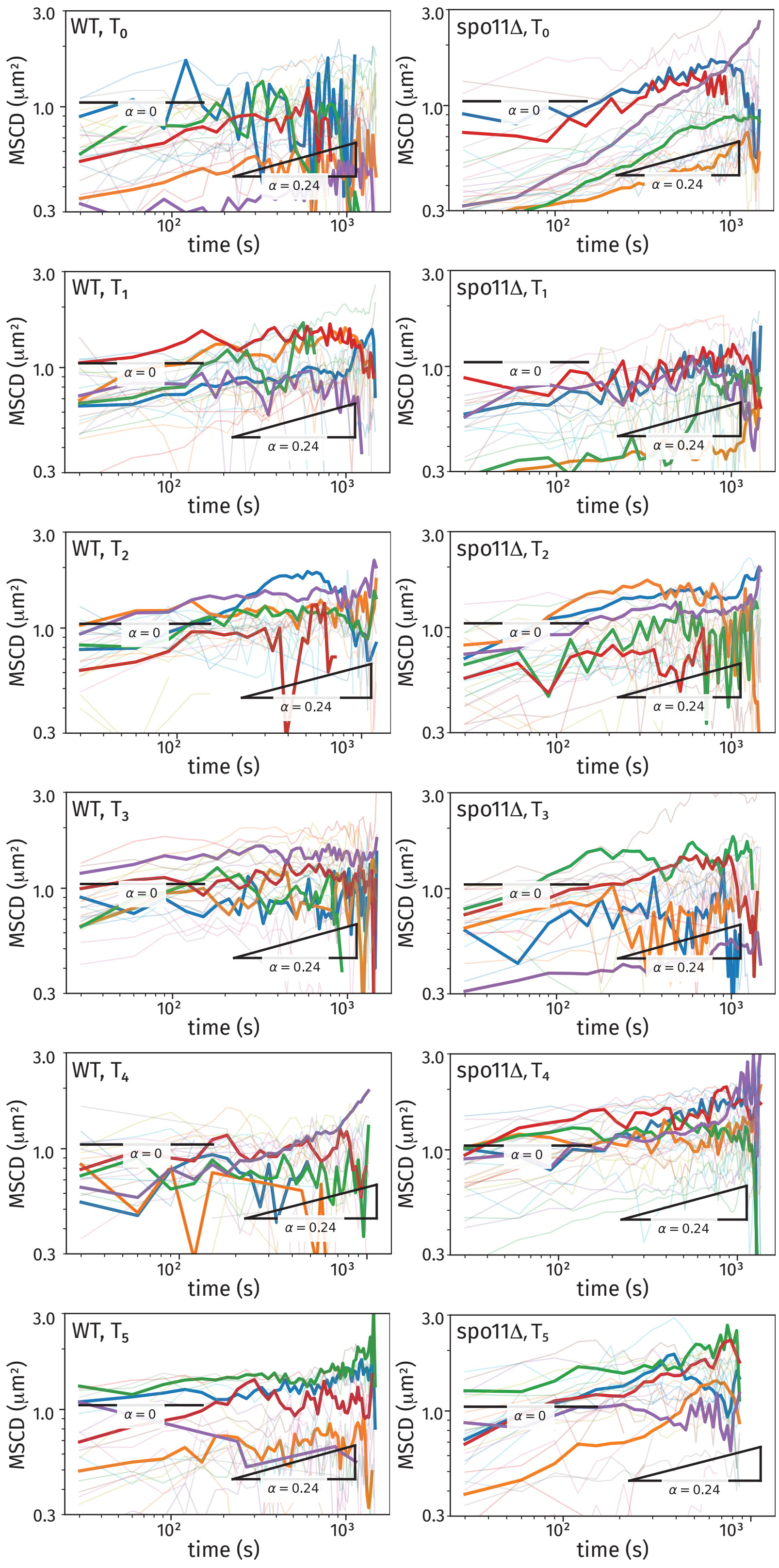


**Fig. S9.** Single-cell MSCD plots for the *LYS2* locus for chronological stage $T_0$ to $T_5$ (same presentation as in Fig. 5 of the manuscript).

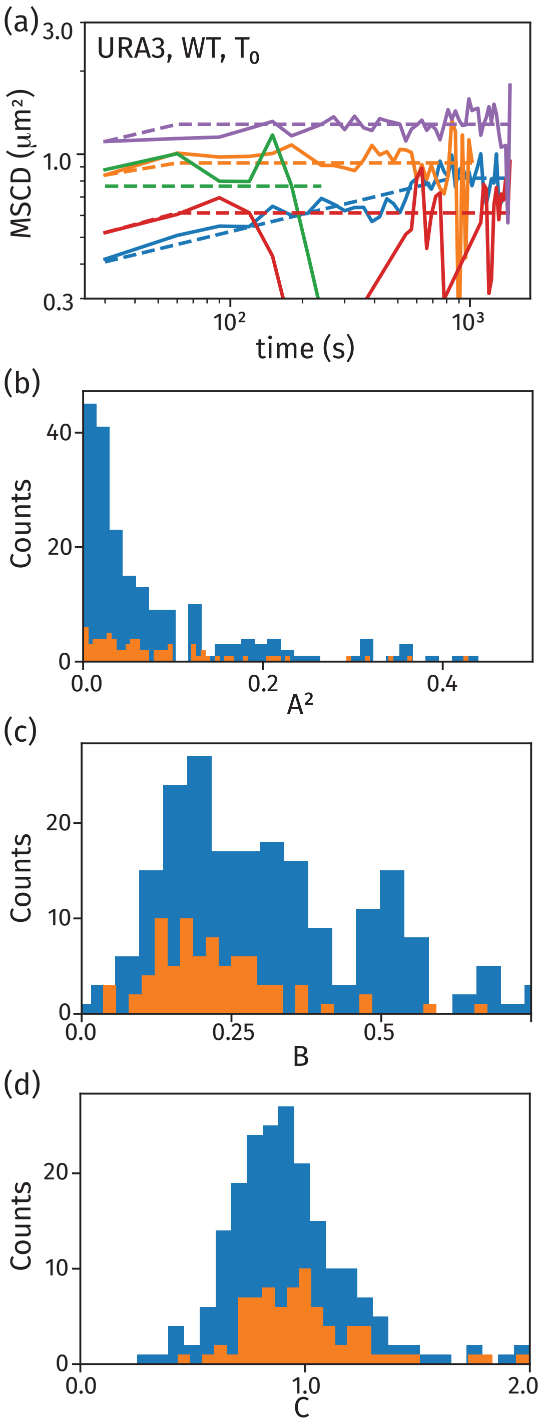


**Fig. S10.** Distribution of model parameters $a$, $b$, and $c$ from fits to the single-cell MSCD with a functional form $MSCD = \min(At^B, C)$ for the *URA3* locus at chromosomal stage $T_0$. The blue histogram shows data from all of the trajectories. The orange histogram is associated with trajectories with over 10 data points of powerlaw behavior before reaching the plateau value.

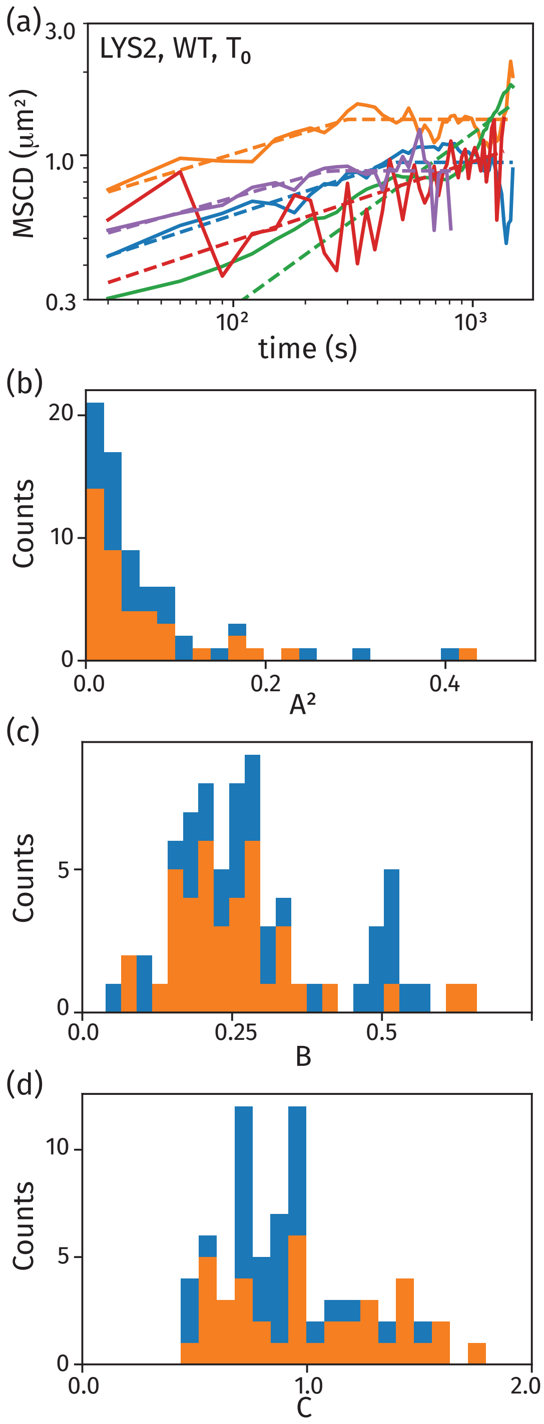


**Fig. S11.** Distribution of model parameters $a$, $b$, and $c$ from fits to the single-cell MSCD with a functional form $MSCD = \min(At^B, C)$ for the *LYS2* locus at chronological stage $T_0$. The blue histogram shows data from all of the trajectories. The orange histogram is associated with trajectories with over 10 data points of powerlaw behavior before reaching the plateau value.

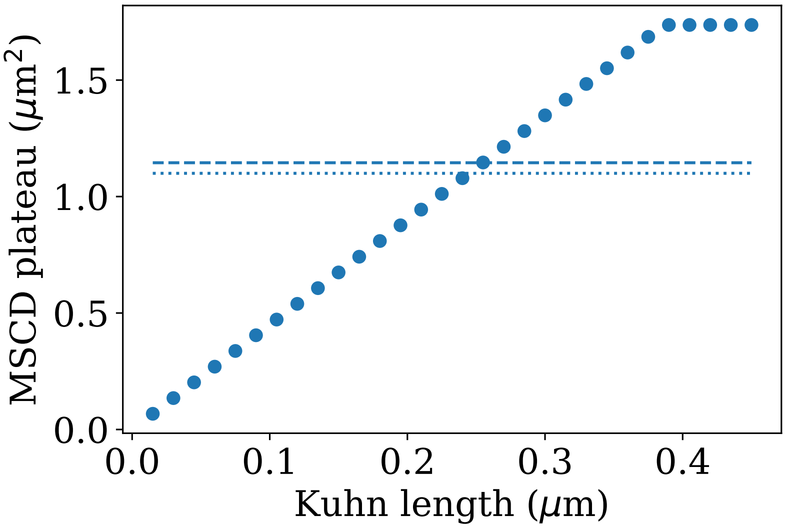


**Fig. S12.** Plot of the MSCD plateau value versus Kuhn length for our model biased on the genomic positions and lengths for the *URA3* locus. The horizontal dashed and dotted lines coincide with the MSCD plateau values at $T_0$ for the wild-type and *spo11*Δ strains of the *URA3* locus, respectively. We apply a linear compaction of 31.6 bp/nm to convert between genomic size (in bp) and chain length (in nm)

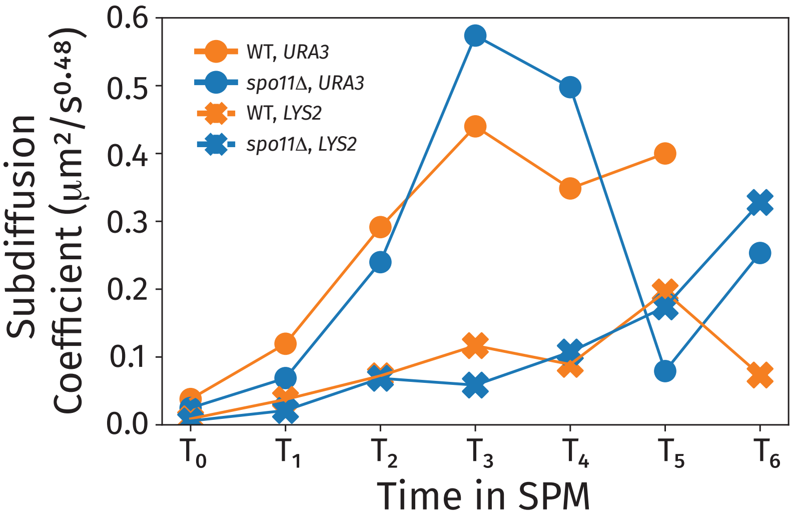


**Fig. S13.** Fitted values of the subdiffusion coefficient versus chronological stage for wild-type and *spo11*Δ strains from experimentally determined ensemble-averaged MSCD of *URA3* and *LYS2* loci.

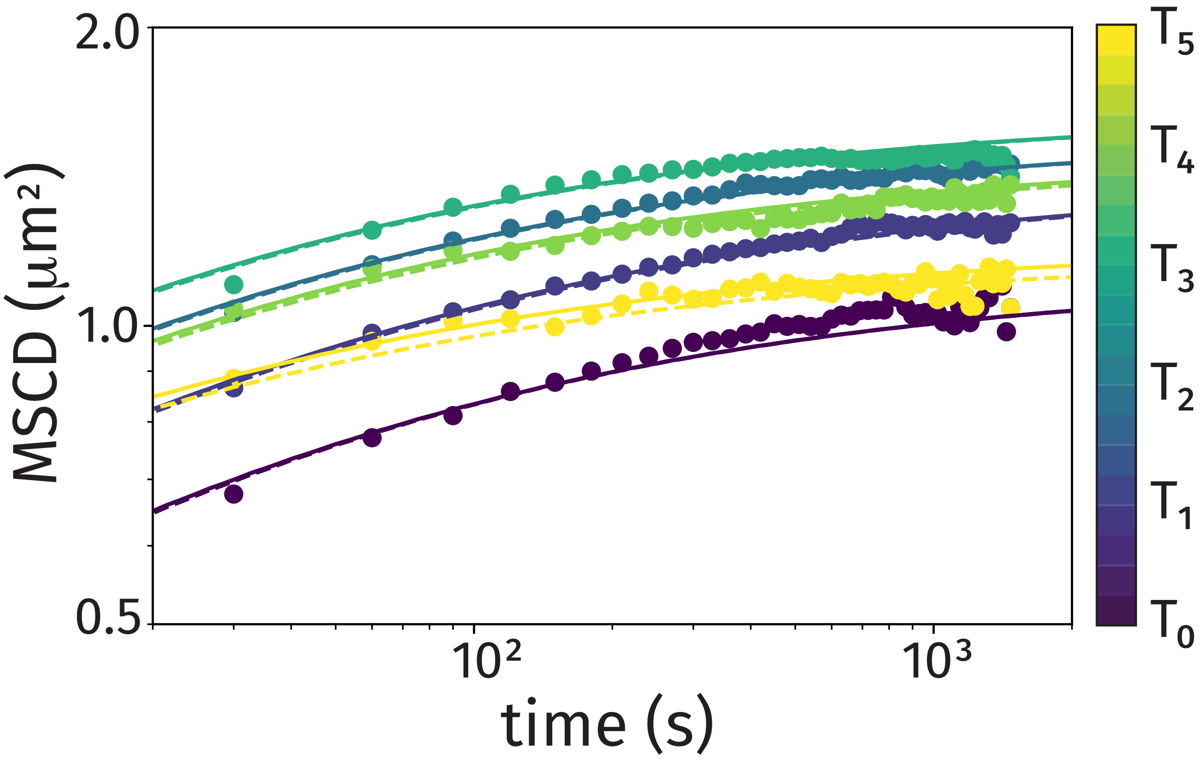


**Fig. S14.** Time-and-ensemble averaged MSCDs at different times after induction of sporulation, for wild-type strain tagged at the *URA3* locus. We include theoretical predictions based on the biased theory (i.e. excluding MSCD values less than 0.0625 $\mu\mathrm{m}^2$ due to imaging resolution) as the solid curves and the unbiased theory (i.e. including all predicted values) as the dashed curves.

| Strain number | Relevant genotype | Source |
|---|---|---|
| SBY2489 | *MATα ho::hisG lys2 leu2::hisg ura3::hisG GAL3* | - |
| SBY3660 | *MATa ho::hisG LEU2::tetR-GFP URA3::tetOx224 his3::hisG spo11::kanMX* | - |
| SBY3662 | *MATa ho::hisG LEU2::tetR-GFP URA3::tetOx224 his3::hisG* | - |
| SBY4110 | *MATa/α leu2::pURA3-tetR-GFP URA3::tetOx224::URA3 ndt80::URA3* | Amon #A6946 |
| SBY4111 | *MATa/α leu2::LEU2-tetR-GFP lys2::tetOx240::URA3 ndt80::URA3* | Amon #A9828 |
| SBY5002 | *MATa ho::hisG LEU2::tetR-GFP URA3::tetOx224 his3::hisG spo11::kanMX GAL3* | This work |
| SBY5004 | *MATα ho::hisG LEU2::tetR-GFP URA3::tetOx224 his3::hisG GAL3* | This work |
| SBY5011 | *MATα ho::LYS2 ura3 leu2::hisG his3::hisG leu2::pURA3-tetR-GFP::LEU2 lys2::tetOx224::URA3 spo11::kanMX* | This work |
| SBY5905 | *MATα ho::hisG LEU2::tetR-GFP URA3::tetOx224 his3::hisG gal3* | This work |
| SBY5907 | *MATa ho::hisG LEU2::tetR-GFP URA3::tetOx224 his3::hisG gal3* | This work |
| SBY5909 | *MATa/α ho::hisG/" LEU2::tetR-GFP/" URA3::tetOx224/" his3::hisG/" gal3/"* | This work |

**Fig. S15.** Yeast strains used in this study. All strains used were in the SK1 background. The original *teto*/TetR-GFP strains were a gift from Angelika Amon.

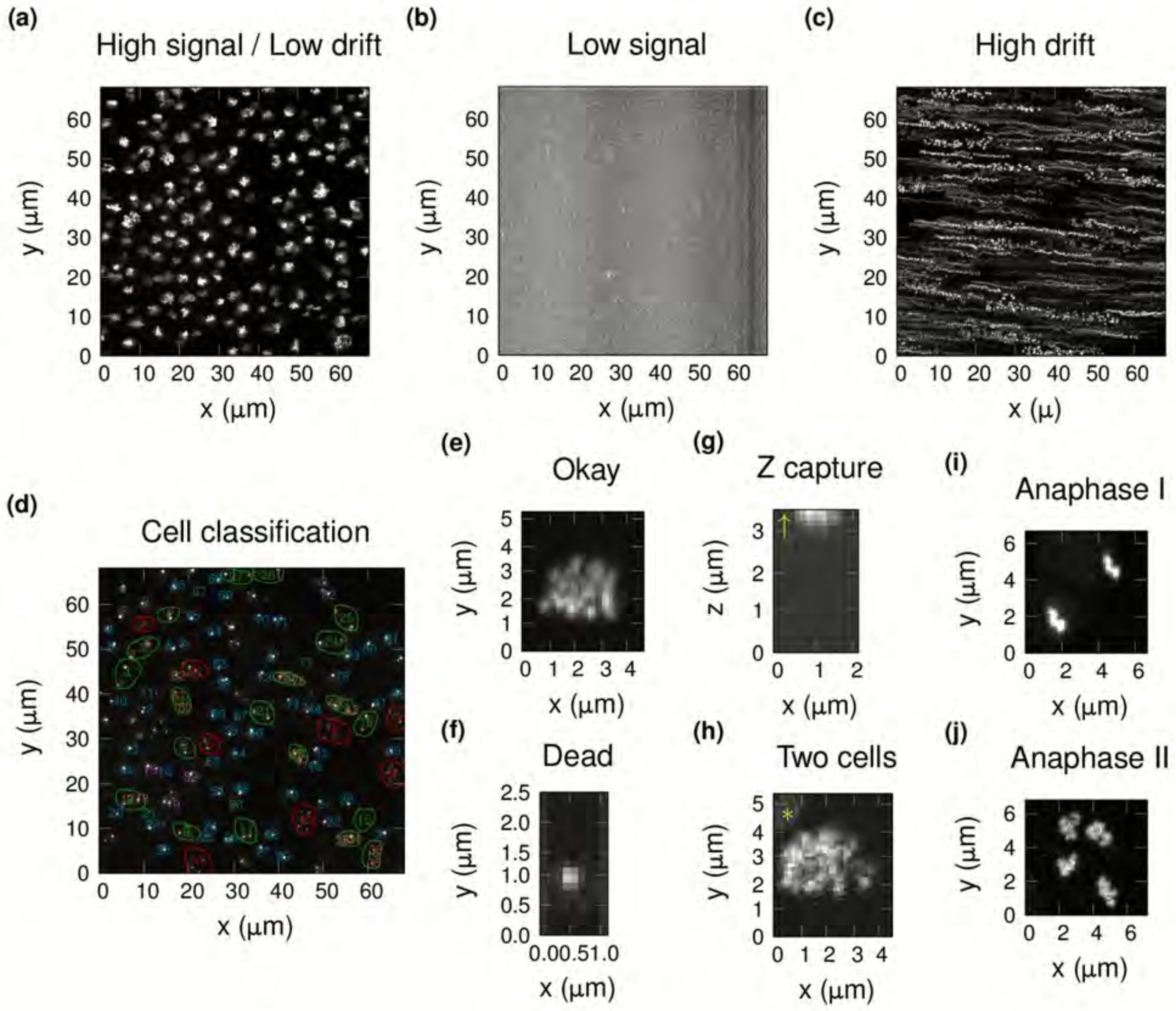


**Fig. S16.** Criteria for assessing time course quality. (a) An example zt-MIP for a quality video with both a high signal to noise ratio and low imaging drift. (b) An example zt-MIP for a video excluded from further analysis due to low signal to noise. (c) An example zt-MIP for a video excluded from further analysis due to high imaging drift. (d) A z-MIP at a single time point, with cropped cells indicated by the cell number, the color of the number reflects an observation made of the cell, e.g. blue is "okay". Cells circled green have progressed past anaphase I, and cells circled red have progressed past anaphase II. Example MIPs are shown for (e) an "okay" cell, (f) a "dead" cell, (g) a cell (yt-MIP) that has drifted out the z-capture range (yellow arrow), (h) a cropped region containing information from two different cells (yellow asterisk), (i) a dyad that has progressed past anaphase I, and (j) a tetrad that has progressed past anaphase II. Only "okay" cells were included in the final dataset.

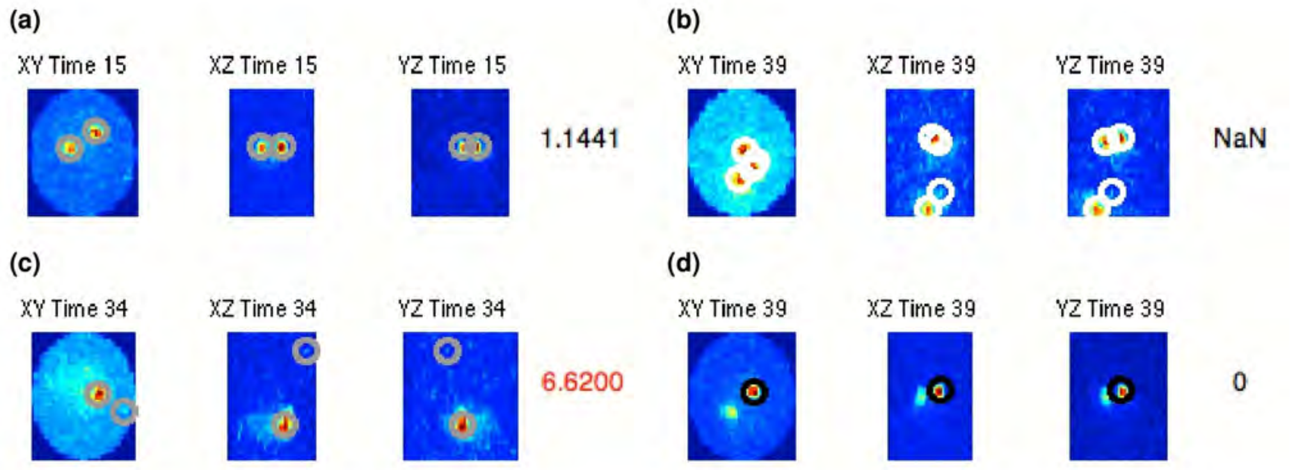


**Fig. S17.** Errors observed when assessing computational spot calling. Single time points are shown for different cells, with three projections shown; z-MIP, y-MIP, and x-MIP. (a) An example of correct spot calling. (b) An example of extra spots, a maximum of two spots could be detected. (c) An example of a type I error (false positive), detecting a second spot when a second spot is not present. (d) An example of a type II error (false negative), failing to detect a second spot when a second spot is present. Time points with spot calling errors were changed to NaN.

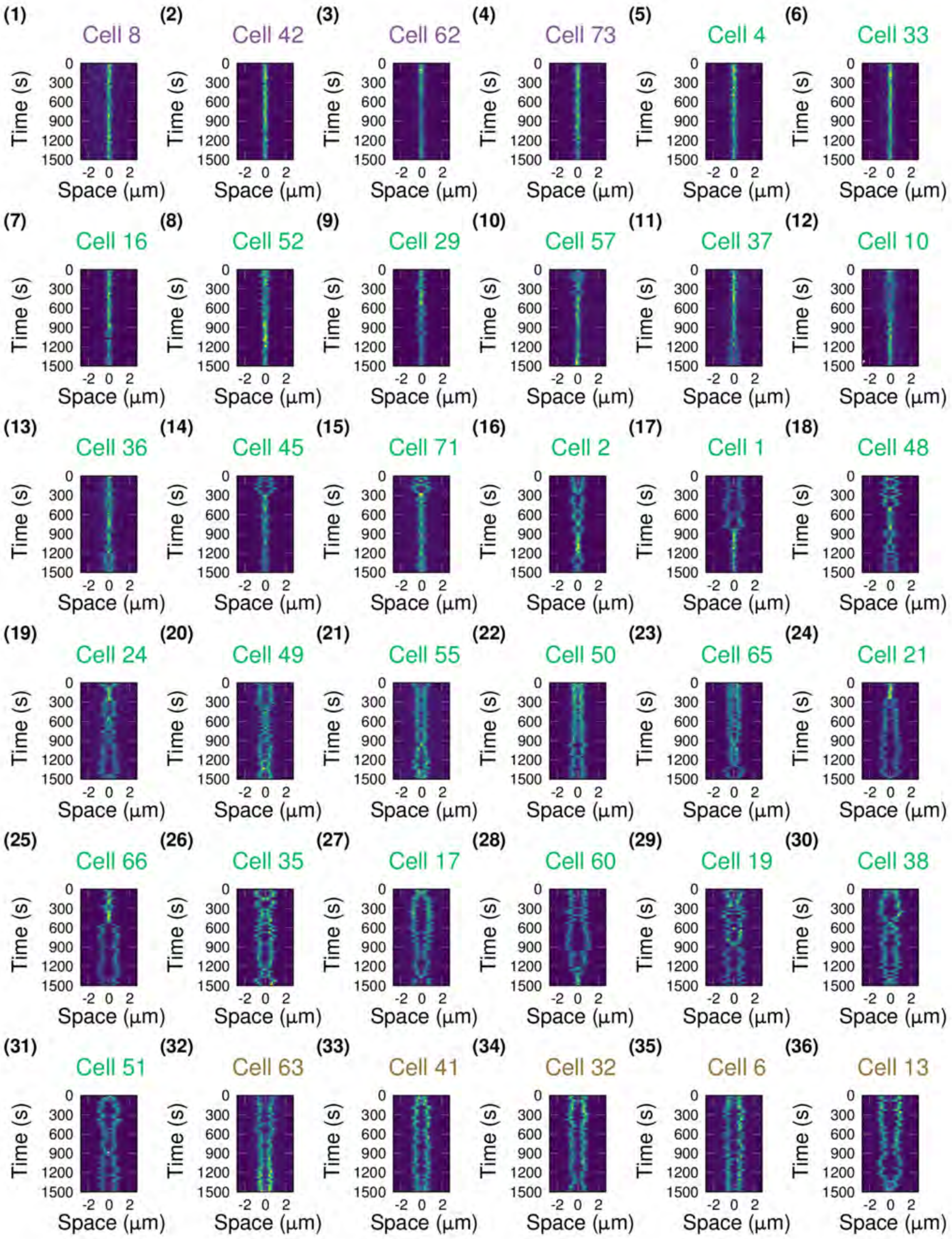


**Fig. S18.** Kymograph gallery for representative wild-type experiment. Kymographs for QC confirmed cells (36/36 are shown) from URA3 wild-type experiment 6, T3. Cells are ordered by the average distance between foci over time. Color of the cell number indicates whether cell was always paired (dark gray), mixed (green), or always unpaired (light gray).

**Supplemental File "Field.avi":** Movie showing live fluorescence captured from a field of cells at T3 showing (a) movement of the spots over time, and the (b) cumulative area explored by foci.

**Supplemental File "Mixed.avi":** Movie showing live fluorescence captured from a cell cropped from a T3 movie, that was "mixed" during observation. Shown is (a) spot position in 3D space over time (4D), and (b) a trace of the distance between the spots over time.

**Supplemental Field "Unpaired.avi":** Movie showing live fluorescence captured from a cell cropped from a T3 movie, that was "always unpaired" during observation. Shown is (a) spot position in 3D space over time (4D), and (b) a trace of the distance between the spots over time.

**Supplemental File "Paired.avi":** Movie showing live fluorescence captured from a cell cropped from a T3 movie, that was "always paired" during observation. Shown is (a) spot position in 3D space over time (4D), and (b) a trace of the distance between the spots over time.

**Supplemental File "supplemental dataset":** XYZ coordinates for foci in all cells over time and across all time courses.
